# Exploring mitonuclear interactions in the regulation of cell physiology: insights from interspecies cybrids

**DOI:** 10.1101/2024.12.03.626354

**Authors:** Kateryna Gaertner, Riikka Tapanainen, Sina Saari, Zsófia Fekete, Steffi Goffart, Jaakko L. O. Pohjoismäki, Eric Dufour

**Affiliations:** Faculty of Medicine and Health Technology, FI-33520 Tampere University, Finland; Department of Environmental and Biological Sciences, FI-80101 University of Eastern Finland, Finland

**Keywords:** mitonuclear interactions, mitochondrial transfer, cytoplasmic hybrid, hares, speciation, metabolism

## Abstract

Brown hares (*Lepus europaeus*) and mountain hares (*Lepus timidus*) frequently hybridize in regions where their range overlaps, producing fertile offspring and enabling gene flow between the species. Despite this, no hybrid species has emerged, suggesting that hybrid backcrosses may incur fitness costs. One potential mechanism for such costs involves the interactions between mitochondrial and nuclear gene products, where incompatibilities between species-specific alleles may reinforce species barriers and lead to hybrid breakdown. However, direct experimental evidence for this hypothesis remains limited.

In this study, we used fibroblasts derived from skin biopsies of wild-caught hares to generate cytoplasmic hybrid (cybrid) cell lines, wherein mitochondria and mtDNA from one species were transferred to mitochondria-depleted cells of the other species, creating novel mitonuclear gene combinations while preserving the original diploid nuclear background. Employing a range of techniques – including transcriptomics, metabolomics, microscopy, and respirometry – we explored the consequences of mitochondrial transfer between these hare species. Our results reveal that in the studied species mitonuclear incompatibilities exhibit strong effects on cellular fitness but are limited to specific genotypes. We propose mechanisms of cellular-level incompatibility and their potential consequences for interspecific hybrids, offering new insights into the complexity of mitonuclear interactions.

## Introduction

Eukaryotic cells represent a unique union of two distinct genetic components. In animals, the biparentally inherited nuclear genome contains on average 18,943 genes [1], whereas the uniparentally inherited mitochondrial DNA (mtDNA) encodes 2 rRNA, 27 tRNA and 13 protein components [2] of the electron transport chain (ETC) complexes and the ATP-synthase, required for ATP production through a process known as oxidative phosphorylation (OXPHOS). While these core OXPHOS components are produced by the mitochondrial genome, the remaining subunits, as well as factors required for the regulation and replication of the mtDNA – are encoded by the nuclear genome. Besides OXPHOS, mitochondria are the central location for numerous vital metabolic and physiological processes [3], including the synthesis of iron-sulphur clusters, regulation of calcium and redox homeostasis; programmed cell death [4]; heat production [5]; detoxification reactions [6], and immunity [7], all of which are driven by nuclear gene products, although directly or indirectly dependent on OXPHOS function.

Due to their reliance on both nuclear and mitochondrial genomes, OXPHOS functions are especially sensitive to genetic incompatibilities, which might also act as a major reinforcer of species boundaries [8]. Initially, the maternal nuclear genome may help maintain compatibility between the maternal mtDNA and the hybrid’s nuclear genome. However, as hybrids backcross with the paternal species, the proportion of the maternal nuclear genome decreases, leading to a loss of this compatibility. Over time, this can result in hybrid breakdown due to mismatches between the maternal mtDNA and the paternal nuclear genome, disrupting essential mitonuclear interactions required for cellular function [9, 10]. Besides imposing selection pressure to purge incompatible mitochondrial genomes in subsequent hybrid lineages, selfish mtDNA haplotypes have been suggested to drive an adaptive arms race between the mitochondrial and nuclear genomes, a phenomenon known as the Red Queen Effect [11]. For example, the mitochondrial haplotype has been reported to influence the evolution of nucleus-encoded mitochondrial genes in highly female philopatric Macaque populations, with wider effects on species diversity, ecology and behaviour [12].

We have been interested in understanding the physiological differences between species and the maintenance of species barrier between two interfertile hare species, the mountain hare (*Lepus timidus*) and the brown hare (*L. europaeus*). Facilitated by the climate change, the temperate-adapted brown hare is expanding its habitat to the North of Europe previously dominated by the cold-adapted mountain hare [13–15]. In this context, hybridizations occur frequently, particularly at the leading edge of the brown hare’s expansion front [16, 17], resulting in biased genetic introgression from the mountain hare to the brown hare [18–22].

Introgression is very common among hares, with many species exhibiting reticulated evolution with frequent hybridization [19, 23, 24], sometimes preserving introgressed mtDNA from locally extinct species in contemporary populations [19, 20]. Interestingly, while the first generation hybrids of female brown hares and male mountain hares are found in nature [18, 21, 25], the subsequent introgression of brown hare mtDNA into the mountain hare populations is very rare [14–17, 22]. This bias may be due to demographic or behavioural factors [17, 19, 20], but it is also possible that mountain hare mtDNA is more compatible with the brown hare nuclear genome than the reciprocal combination.

Compared to mountain hares, brown hares invest more in growth and reproduction at the expense of longevity [26]. Given the key role of mitochondria in metabolism and aging, it is possible that nuclear-mitochondrial coevolution has been more stringent in mountain hares, for example due to the need for energy-efficient adaptations to survive harsh winters with limited or poor-quality food sources. These appear to be reflected at the cellular level, as brown hare fibroblasts have greater proliferative capacity and higher metabolic rate than mountain hare cells [27], features likely reflecting differences in the species’ life history strategies.

To test the effect of introgressed mitochondrial DNA on cell physiology and to identify potential incompatibilities between the nuclear and mitochondrial genomes of the two hare species, we generated cytoplasmic hybrid (cybrid) cell lines combining mtDNA from one species with the nuclear genome of the other. A similar approach has been previously used to study mtDNA mutations in human patients [28, 29] and interspecific incompatibility between nuclear and mitochondrial OXPHOS components [30]. We here applied this method for the first time in combination with detailed analyses of cell physiology, metabolomics, and transcriptomics to uncover the broad-scale effects of disrupted mitonuclear crosstalk. We find that while cybrid cells retain most features of the nuclear host, the process of cybrid generation induces specific changes in gene expression and metabolism that are independent of the mitochondrial haplotype. Generally, cybrid cells exhibit intermediate phenotypes for many traits compared to the host cells. However, cells with a mountain hare nucleus and brown hare mtDNA displayed more pronounced, though inconsistent, changes relative to their reciprocal cybrids, which showed signs of an overall increase in metabolic activity. Notably, the cybrids also displayed a distinct metabolic profile from that of the mitochondrial donor cells. Our results offer insights into altered mitonuclear interactions in species hybrids, while also highlighting the need for caution when interpreting cybrid-based studies of human mitochondrial disorders.

## Materials and methods

### Isolation and culture of hare cells

Four mountain hare (LT1, LT4, LT5, LT6) and four brown hare (LE1, LE2, LE3, LE4) fibroblast cell lines were established from skin biopsies [27] and maintained in standard cell culture environment at 37 °C and 5 % CO_2_ in high glucose (4.5 g/l) DMEM, supplemented with 10 % FBS, 1 % L-glutamine (Sigma-Aldrich) and 1 % P/S (Gibco). In addition to the standard cell lines, mtDNA-less ρ0-cells were generated by culturing the cells in the presence of 100 μM ddC, until no mtDNA was detected on a Southern blot [31]. The ρ0-cells were maintained in high glucose DMEM with 10 % FBS and 50 μg/ml uridine to complement the loss of mitochondrial functionality on pyrimidine biosynthesis [32].

### Generation of the cybrid cell lines

The cybrid cells were generated by fusing a ρ0 host cell with an mtDNA donor, whose nucleus had been destroyed by Actinomycin D treatment [33]. Three reciprocal pairs of heterocybrid cell lines were generated, taking advantage of sex and haplotype differences to control for the nuclear and mitochondrial identities: LE1♂ x LT4♀, LE2♀ x LT1♂ and LE3♀ x LT6♂. In addition, two control homocybrid cell lines with mtDNA from the same species but a different individual were generated: LE1♂(nucleus) x LE2♀ (mtDNA) and LT6♂ (nucleus) x LT4♀ (mtDNA). Each cybrid cell line was subcloned on 96-well plates to ensure their genetic uniformity. Please see [31] for the methodological details. For clarity, we refer to the cells providing the nucleus as the “parental” cells and those providing the mitochondria as the “mitochondrial donors”.

### mtDNA sequence difference comparisons

The complete annotated mitochondrial genomes of the hare cell lines used in this study have been published earlier [31]. The polypeptide sequences for the mitochondrial OXPHOS complexes I (ND1, ND2, ND3, ND4, ND4L, ND5, ND6), III (CYTB), IV (COX1, COX2, COX3) and V (ATP6, ATP8) were aligned using Geneious 10.2.6 to calculate their amino acid differences.

### Cell morphology measurements

Live cells grown on glass-bottom dishes (MatTek) were stained with CellMask Actin (Act), nuclei with NucBlue (NB), and mitochondria with 100 nM TMRM (Invitrogen) in the presence of 50 µM Verapamil (Sigma-Aldrich) [34], an efflux pump inhibitor [35]. To promote optimum cell health throughout the imaging procedure, the cells were maintained in a stage-top incubator (Tokai Hit) pre-set to standard culture conditions in the presence of FluoroBrite DMEM and 1 % P/S (Gibco). Relative z-stacks were acquired from ∼5 non-overlapping imaging fields using a 40x / 1.15 water-immersion objective and Perfect Focus System of A1R+ confocal microscope system (Nikon Eclipse Ti2-E). Detector settings for NB (Ex 405, Em 450/50), Act (Ex 488, Em 525/50) and TMRM (Ex 561, Em 595/50) were independently adjusted to avoid image saturation. Images were deconvolved with Huygens Essential (SVI; Hilversum) and analyzed in Fiji-ImageJ [36]. Maximum intensity z-projections were generated from the Act channel to measure cell size with the Polygon Selection tool and from the NB channel to measure nucleus size with the Magic Wand tool (n = 40 images / cell line). Midsections of individual cells were used to measure mitochondrial morphology with the Mitochondria Analyzer plugin as previously described [37, 38]. Shapes of nuclei and mitochondria were compared using the aspect ratio that ranges from 0 “round” to 1 ≥ “elongated”. Mitochondrial and nuclear volumes were defined as a relative size of the organelle in a given cell: 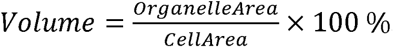.

### Cell growth and metabolic activity assays

Cells were seeded in sextuplicate at densities of 1×10^4^ cells/well into 48-well plates corresponding to 8 time points. General metabolic activity was measured with MTT assay as described [39]. Briefly, the cells in three wells were incubated in the presence of 0.5 mg/ml MTT for 3 h at standard culture conditions, and the resulted formazan was dissolved in DMSO (Sigma-Aldrich). Three other wells from the same plate were used to estimate cell growth with sulforhodamine B (SRB) assay [40]. Briefly, cells were fixed in 4 % paraformaldehyde (PFA) and stained with 0.4 % SRB (Sigma-Aldrich) diluted in 1 % acetic acid for 30 min at room temperature (RT), washed and solubilized in 10 mM Tris base (pH 10.5). The absorbance of the resulting solutions of MTT (λ = 570 nm) and SRB (λ = 564 nm) were measured with a Spark microplate reader (Tecan). Blank wells containing no cells were processed identically to define backgrounds of the staining methods. Background-subtracted absorbance (i.e., corrected absorbance) values were used for statistical analyses. The relative cell number measured by SRB and the metabolic activity measured by MTT were plotted (*y*) against time (*x*) to obtain growth and viability curves. A linear regression model was applied to the log phase of cell growth (i.e., days 3 - 7), and comparison between slopes was made using one-way ANOVA with Tukey’s correction in R 4.3.2.

### Wound closure rate measurements

Th wound healing assay was done in triplicate as previously described [27, 41]. Briefly, a manual scratch was performed with a sterile micropipette tip through a confluent cell monolayer. Cells were rinsed with medium and placed in a stage top incubator (Okolab) adjusted to maintain standard culture conditions. Phase contrast images were captured from three non-overlapping fields using 5x / 0.12 dry objective of DMi8 microscope (Leica) every 1 h during 43 h. Time-lapse images were analyzed with the MRI Wound Healing tool [42] in Fiji-ImageJ [36]. Wound closure rate (WCR) was defined as an average velocity at which cells collectively move into the gap, and it was calculated from slopes of the linear (R^2^ ≥ 98%) cell migration phase: 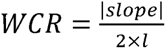, where *l* is the gap length (Jonkman et al. 2014).

### ROS and mitochondrial membrane potential measurements

For reactive oxygen species (ROS) measurements, cells were collected by trypsinization in quadruplicate, pelleted at 250 g for 3 min unless otherwise stated, resuspended with DPBS at 1×10^6^ cells/ml and split into 2 groups: 1) untreated cells; 2) treated with 0.5 µM Rotenone (Sigma-Aldrich), an inhibitor of complex I that drives ROS production [43]. The samples, excluding negative controls, were stained for 30 min at 37 °C with 5 µM CellROX Green (Invitrogen) that exhibits fluorescence upon oxidation by ROS, washed with DPBS, pelleted and resuspended in fluorescence-activated cell sorting (FACS) buffer consisting of DPBS and 2 % FBS. Mean fluorescent intensities of CellROX Green (Ex 488 nm, Em 525/40 nm) were measured by recording 40 000 events/sample at medium speed with CytoFlex S flow cytometer (Beckman Coulter). Mean signal intensities from unstained samples (negative controls) were subtracted from CellROX stained samples, and the resulting values were used for statistical analyses. For visualization purposes, cells were stained with NB and CellROX Green and imaged as described for cell morphology measurements.

For mitochondrial membrane potential (Δψm) measurements, the collected cells were split into 4 groups: 1) untreated cells; 2) treated with 0.5 µM Rotenone; 3) treated with 1.5 µM Oligomycin, a mitochondrial ATP synthase inhibitor; 4) treated with 2 µM FCCP (Sigma-Aldrich), an uncoupler of the electron transport chain. Samples, excluding negative controls, were stained at 37 °C for 20 min with 20 nM TMRM (Invitrogen), a Δψm-sensitive probe, in the presence of 50 µM Verapamil (Sigma-Aldrich), washed and resuspended in FACS buffer. Median fluorescent intensities of TMRM (Ex 561 nm, Em 584/42 nm) were measured as described above by flow cytometer. All data were analyzed with BD FlowJo 10.7.2 software.

### Biolog phenotype assay

Cells were seeded at 1.5×10^4^ cells/well into carbon (PM-M1) and nitrogen (PM-M2) substrate MicroArrays (Biolog) and cultured for 40 h in the presence of IF-M2 medium (Biolog) supplemented with 5% FBS, 1% P/S and either 0.3 mM L-glutamine (PM-M1) or 2 mM galactose (PM-M2). A redox MB dye (10 µl/well, Biolog) was added and incubated with cells for additional 24 h. Absorbance from the accumulated signal (NADH, λ = 590 nm) was measured at 37 °C with a Spark microplate reader (Tecan). Wells containing cells with no substrates (background wells) were processed identically to define NADH signal backgrounds for every cell line and subtracted from values of substrate containing wells (i.e., corrected absorbance). This approach allowed to categorize substrates into 3 groups: “-” disliked substrates that reduced cell growth or caused cell death, “∼ 0” neutral substrates that did not substantially differ from wells containing no substrates (background wells), “+” preferred substrates that were utilized by cells and supported their growth. Two independent experiments were performed for each cell line, except for LE2 (nucleus) x LT1 (mtDNA) heterocybrid, where the cells died in one of the experiments. Corrected mean values were used for statistical analysis (two-way ANOVA with Tukey’s correction). For heatmap [44] generation, hierarchical clustering of cell lines (columns) was performed using Euclidean distance and the complete linkage method. Rows (substrates) were grouped into three categories (disliked, neutral, preferred) using k-means partitioning, implemented in R version 4.3.2.

### Cell respiration

High-resolution respirometry was conducted in septuplicate using Oxytherm (Hansatech instruments) as previously described [27]. Basal respiration of digitonin-permeabilized cells was measured by recording oxygen consumption rates (OCR) after stimulation of OXPHOS with 5 mM Pyruvate, 5 mM Glutamate, 5 mM Malate, 1 mM ADP (Complex I respiration) and 5 mM Succinate (Complex I+II respiration). ATP-coupled respiration was measured after inhibition of Complex V with 1 µM Oligomycin. Maximal respiration was evaluated by uncoupling with 1 µM FCCP, after which the respiration from Complex I was blocked with 0.3 µM Rotenone, and from Complex III with 90ng/ml Antimycin A (Maximum capacity). The proton leak was measured as a difference between Complex V-blocked respiration (i.e., injection of Oligomycin) and Complexes I- and III-blocked respiration (i.e., injection of Rotenone and Antimycin A). Spare respiratory capacity was defined as the difference between the maximal respiration and the basal respiration. Treatment with 700 µM Ascorbate and 300 µM TMPD, an artificial electron donor [45], resulted in maximal Complex IV respiration that was completely blocked by 0.2 mM KCN (Sigma-Aldrich).

Respirometry of intact cells (n = 5–8 per cell line) was measured with a Seahorse Analyzer (Agilent) following the manufacturer’s guidelines. Briefly, cells were seeded at 2.5×10^4^ cells/well into XF24 plates (Agilent) 22 h prior to the assay. Cells were washed and incubated at 37°C for 1 h in a non-CO_2_ incubator with an assay medium consisting of Seahorse XF DMEM (pH 7.4) supplemented with 10 mM glucose, 1□mM Pyruvate and 2□mM L-glutamine (Agilent). Mitochondrial respiration was assessed with a Cell Mito Stress Assay, and bioenergetic profiles were measured with Real-Time ATP Rate Assay and Long Chain Fatty Acid (LCFA) Oxidation Stress Assay (Agilent). Concentrations of the used inhibitors were: 1.5 μM Oligomycin, 2□μM FCCP, 0.5□μM Rotenone, 0.5□μM Antimycin A, and 4 μM Etomoxir, an inhibitor of carnitine palmitoyltransferase 1 (CPT1) that interrupts the fatty acid oxidation (FAO) pathway [46]. Data analyses were done using Seahorse Analytics 1.0.0-699 (Agilent).

### Targeted metabolomics

Cybrids were seeded (n = 6 per cell line) and grown for 3 days until they reached 80 % confluence. The medium was changed 4 h prior metabolite extraction. Cells were rinsed with ice-cold 150 mM NH4AcO (pH 7.3) and incubated with 1 ml/well 80 % MeOH at −80 °C for 45 min. Cells were scraped and centrifuged at 1.6×10^4^ g for 15 min at 4 °C. Pellets were lysed in 0.1 M NaOH and used for sample normalization by protein concentrations estimated with BCA assay kit (Thermo Scientific). Supernatants were dried using a vacuum concentrator (ScanVac MaxiVac, LaboGene) and stored at −80 °C. Samples were sent for targeted analysis to the FIMM Metabolomics Unit at the University of Helsinki (Helsinki, Finland). The metabolites were detected with a Thermo Vanquish UHPLC coupled with Q-Exactive Orbitrap quadrupole mass spectrometer (MS) equipped with a heated electrospray ionization (H-ESI) source probe (Thermo Fischer Scientific) using full MS and polarity switching mode in mass range 55 to 825 m⁄z and resolution of 70.000. Samples were delivered in acetonitrile:methanol:Milli-Q water (40:40:20) and the chromatographic separation was done using SeQuant ZIC-pHILIC (2.1×100 mm, 5μm particle) column (Merck) at a flow rate of 100 µl/min. TraceFinder 4.1 software (Thermo Fischer Scientific) was used for data integration, the final peak integration and peak area calculation of each metabolite. Four metabolites with less than 50 % of non-zero measures per group and three samples (3x LE2 (nucleus) x LT1 (mtDNA)) were excluded as significant outliers. Metabolite levels were normalized by protein content. The data were analysed using MetaboAnalyst 6.0 [47] and GraphPad Prism 9.5.1.

### Transcriptome analysis

RNA isolation, mRNA purification has been described previously [27]. RNA sequencing quality control and trimming of adapters and low-quality sequences was done using fastp [48], with the options --length_required 50 and –detect_adapter_for_pe. Alignment to the *Lepus europaeus* reference genome (GenBank accession GCA_033115175.1) [49] was done with the default options of STAR 2.7.11. [50]. Variant calling was done by using bcftools mpileup and call [51, 52], with default parameters. The called variants were filtered based on overall call quality (QUAL > 100), mapping quality (MQ > 20), mapping quality bias (−1< MQBZ < 1), base quality bias (−1 < BQBZ < 1), minimum read depth (DP > 10) and for mapping quality *vs* strand bias (−1 < MQSBZ < 1), resulting in 214,641 variants after filtering. Genotype matrix was generated from the vcf file with a Python script, the eigenvectors of the PCA were calculated with the prcomp function in R [53]. For the differential expression analysis, reads were quantified by htseq-count [54].

From all annotated genes (30,833), the ones having ≤ 20 reads across all 15 cell lines were filtered out. Differential expression analysis of the remaining 19,357 genes was performed using DESeq2 1.42.1 [55]. Benjamini-Hochberg method [56] was used to adjust *p*-values. First, to remove potential bias resulted from the cybrid generation process, control homocybrids were grouped together with heterocybrid cell lines based on their nucleus background (with either LE or LT) and contrasted with the respective parental cell lines (i.e., all LE cybrids *vs* LE, all LT cybrids *vs* LT). Commonly upregulated or downregulated based on log2 fold change (log2FC) differentially expressed genes (DEGs) were defined as “likely confounding genes” (284, among which 73 are human orthologs) shared between all cybrids regardless of their mtDNA or nucleus origin. Next, to evaluate the effect of introduced foreign mtDNA (interspecies mitonuclear compatibility), control homocybrids were grouped together with parental cell lines (either LE or LT) and contrasted with heterocybrids based on their nucleus background (i.e., LE homocybrid and LE *vs* LE (nucleus) x LT (mtDNA) heterocybrids; LT homocybrid and LT *vs* LT (nucleus) x LE (mtDNA) heterocybrids). Common confounding genes were filtered out, which resulted in species-specific heterocybrid DEGs (295 for LE (nucleus) x LT (mtDNA) heterocybrids and 123 for LT (nucleus) x LE (mtDNA) heterocybrids). Based on z-scores and heatmap analysis [44], we identified “high confidence” heterocybrid DEGs that were different between heterocybrids and control homocybrid and considered them particularly triggered by interspecies mitonuclear interactions, while other DEGs found similar between control homocybrid and heterocybrids were considered as “species-specific confounding genes” that could resulted from species-specific reaction on cybrid generation process or from species-specific interindividual incompatibilities. Gene functional information of shared with human orthologs was retrieved using NCBI, UniProtKB/Swiss-Prot as well as literature search with PubMed.

### Gene expression change analysis

For each heterocybrid cell line, the expression of each gene was compared to that of its two parents, leading to 10 comparisons: LE2 (nucleus) x LT1 (mtDNA) *vs* LE2 or *vs* LT1; LT1 (nucleus) LE2 (mtDNA) *vs* LE2 or *vs* LT1; LE3 (nucleus) x LT6 (mtDNA) *vs* LE3 or *vs* LT6; LT6 (nucleus) x LE3 (mtDNA) *vs* LE3 or *vs* LT6; LE1 (nucleus) x LT4 (mtDNA) *vs* LE1 or *vs* LT4. Based on the direction of the change (increase or decrease), each gene was assigned to one of the 2^10^ = 1024 possible combination of changes in expression. A Monte Carlo multinomial test (10^7^ iterations) was performed to identify any deviation from random distribution of the gene into the 1024 categories. Bonferroni-corrected *post-hoc* goodness of fit Chi^2^ test were performed separately for each of the 1024 category. Monte Carlo multinomial test was performed with and without filtering change in expression from 0 (no filtering) to 100 % (minimum of 2x change) with 10 % steps. The categories of interest were defined as significantly different from random distribution at each level of filtering.

### Mitochondrial DNA copy number analysis

The mtDNA copy number was measured using Real-Time qPCR with the following primers and probes targeting sequences identical in both hare species:

mtDNA hare qPCR F: 5′-ACC CCG CCT GTT TAC CAA-3′

mtDNA hare qPCR R: 5′-ATG CTA CCT TTG CAC GGT CA-3′

mtDNA hare probe: 5′-FAM-TGC CTG CCC AGT GAC AAA CGT-3′

SDH hare qPCR F: 5′-CCT GCC TGG CAT TTC TGA GA-3′

SDH hare qPCR R: 5′-ATT GGC TCC TTG GTG ACG TC-3′

SDH hare probe: 5′-HEX-GCC ATG ATC TTC GCG GGT GTG-3′

The primers were first tested on LE and LT samples using a PCR program with a 3-minute denaturation at 95 °C, followed by 35 cycles of 95 °C for 15 seconds, 60 °C for 15 seconds, and 72 °C for 15 seconds, and a final elongation at 72 °C for 5 minutes (AccuStart II™ PCR SuperMix, Quantabio). Strong PCR products were observed for both species on a 1.5 % TAE agarose gel. For qPCR analysis, the program included a 3-minute denaturation at 95 °C, followed by 40 cycles of 95 °C for 15 seconds, 54 °C for 15 seconds, and 72 °C for 15 seconds. The same master mix was used with fluorescent probes. Each sample was run in triplicate, with standards of 100, 25, 10, 2.5, and 1 ng/µl.

### Western blotting

Protein samples (n = 4 per cell line) were prepared as previously described [27]. Samples (30 μg per sample) were loaded in TGX Stain-Free Protein Gels with one lane dedicated to a protein standard (Bio-Rad) and run at 100 V. Separated proteins were transferred using a Trans Blot Turbo system on methanol-activated LF-PVDF membranes (Bio-Rad). Protein transfer was assessed following instructions for Stain-Free technology (Bio-Rad). Membranes were blocked in EveryBlot Blocking Buffer (Bio-Rad) and probed with either OXPHOS antibody cocktail for CI-ND6, CIII-UQCRC2, COX IV-2, CV-ATP5A (# 45-8199,

Invitrogen, 1:1000) or VDAC1 (# SAB5201374, Sigma-Aldrich, 1:1000) at 4 °C overnight. Membranes were washed and incubated with fluorophore-conjugated secondary antibodies (# 12004159, Bio-Rad) following the manufacturer’s instructions, and imaged with ChemiDoc MP Imaging System (Bio-Rad) using optimal auto-exposure settings for each protein of interest. Densitometric analysis was performed using total protein normalization (TPN) in Image Lab 6.0.1 (Bio-Rad).

### Statistical analysis

Statistical testing was conducted for all measured traits in cell line comparisons. Outliers were identified and removed using ROUT (FDR 1 %) method. Lognormal *vs* normal data distributions were assessed based on normality tests (i.e., Anderson-Darling, D’Agostino, Shapiro-Wilk and Kolmogorov-Smirnov), a maximum likelihood method and generated QQ plots. If lognormal distribution was more likely, the data were log2 transformed and statistics were performed on logs. Unpaired two-tailed t-test or one-way ANOVA with Tukey’s correction for multiple testing were applied, *p* adj. value < 0.05 and, for gene expression data, −1 > log2FC > 1 were considered as statistically significant. Analyses and graphical representations of the obtained results were done in GraphPad Prism 9.0.0, if not stated otherwise.

## Results

We attempted to generate three reciprocal pairs of heterocybrids, as well as two homocybrid cell lines as controls (Table 1). Pairs were selected so that the host cell and mitochondrial donor were of different sexes, allowing to control the origin of the cybrid nuclear and mitochondrial genomes using genotyping and PCR-Restriction Fragment Length Polymorphism (RFLP), respectively [25]. We previously sequenced the complete mitochondrial genomes of the cell lines [31], revealing that coding differences between mountain hare and brown hare mtDNAs were most pronounced in the genes encoding Complex I and Complex V subunits, with approximately 3% variation in their polypeptide sequences (Table 2). Notably, no intraspecific variation was observed among brown hare mtDNAs, whereas all three mountain hare samples had sequence polymorphisms across all Complex I subunits.

**Table 1.** Names and genotypic identities of the parental and cybrid cell lines. Brown hare (*Lepus europaeus*) abbreviated as LE and mountain hare (*L. timidus*) as LT.

| Cell line name | Type | Nucleus (host) | Mitochondria (donor) | Sex | mtDNA haplotype |
| --- | --- | --- | --- | --- | --- |
| LE1 | Parental | LE1 | LE1 | Male | *europaeus* |
| LE2 | Parental | LE2 | LE2 | Female | *europaeus* |
| LE3 | Parental | LE3 | LE3 | Female | *europaeus* |
| LT1 | Parental | LT1 | LT1 | Male | *timidus* |
| LT4 | Parental | LT4 | LT4 | Female | *timidus* |
| LT6 | Parental | LT6 | LT6 | Male | *timidus* |
| LE1n+LT4m | Heterocybrid | LE1 | LT4 | Male | *timidus* |
| LT4n+LE1m | Heterocybrid | LT4 | LE1 | Female | *europaeus* |
| LE2n+LT1m | Heterocybrid | LE2 | LT1 | Female | *timidus* |
| LT1n+LE2m | Heterocybrid | LT1 | LE2 | Male | *europaeus* |
| LE3n+LT6m | Heterocybrid | LE3 | LT6 | Female | *timidus* |
| LT6n+LE3m | Heterocybrid | LT6 | LE3 | Male | *europaeus* |
| LE1n+LE2m | Homocybrid | LE1 | LE2 | Male | *europaeus* |
| LT6n+LT4m | Homocybrid | LT6 | LT4 | Male | *timidus* |

**Table 2.** Polypeptide sequence identities in mtDNA-encoded subunits of Complex (C) I, III, IV and V between mountain hare and brown hare cell lines. Note that all Complex II subunits are encoded by the nuclear genome and therefore not included.

**CI (NDs)**
|  | LE1 | LE2 | LE3 | LT1 | LT4 | LT6 |
| --- | --- | --- | --- | --- | --- | --- |
| LE1 |  | 100 % | 100 % | 97.9 % | 98.0 % | 98.01 % |
| LE2 | 100 % |  | 100 % | 97.9 % | 98.0 % | 98.01 % |
| LE3 | 100 % | 100 % |  | 97.9 % | 98.0 % | 98.01 % |
| LT1 | 97.9 % | 97.9 % | 97.9 % |  | 99.9 % | 99.81 % |
| LT4 | 98.0 % | 98.0 % | 98.0 % | 99.9 % |  | 99.91 % |
| LT6 | 98.0 % | 98.0 % | 98.0 % | 99.8 % | 99.9 % |  |

**CIII (CYTB)**
|  | LE1 | LE2 | LE3 | LT1 | LT4 | LT6 |
| --- | --- | --- | --- | --- | --- | --- |
| LE1 |  | 100 % | 100 % | 98.95 % | 98.95 % | 98.95 % |
| LE2 | 100 % |  | 100 % | 98.95 % | 98.95 % | 98.95 % |
| LE3 | 100 % | 100 % |  | 98.95 % | 98.95 % | 98.95 % |
| LT1 | 98.95 % | 98.95 % | 98.95 % |  | 100 % | 100 % |
| LT4 | 98.95 % | 98.95 % | 98.95 % | 100 % |  | 100 % |
| LT6 | 98.95 % | 98.95 % | 98.95 % | 100 % | 100 % |  |

**CIV (COXs)**
|  | LE1 | LE2 | LE3 | LT1 | LT4 | LT6 |
| --- | --- | --- | --- | --- | --- | --- |
| LE1 |  | 100 % | 100 % | 99.9 % | 99.9 % | 99.8 % |
| LE2 | 100 % |  | 100 % | 99.9 % | 99.9 % | 99.8 % |
| LE3 | 100 % | 100 % |  | 99.9 % | 99.9 % | 99.8 % |
| LT1 | 99.9 % | 99.9 % | 99.9 % |  | 100 % | 99.9 % |
| LT4 | 99.9 % | 99.9 % | 99.9 % | 100 % |  | 99.9 % |
| LT6 | 99.8 % | 99.8 % | 99.8 % | 99.9 % | 99.9 % |  |

CV (ATPs)
|  | LE1 | LE2 | LE3 | LT1 | LT4 | LT6 |
| --- | --- | --- | --- | --- | --- | --- |
| LE1 |  | 100 % | 100 % | 96.9 % | 97.3 % | 97.3 % |
| LE2 | 100 % |  | 100 % | 96.9 % | 97.3 % | 97.3 % |
| LE3 | 100 % | 100 % |  | 96.9 % | 97.3 % | 97.3 % |
| LT1 | 96.9 % | 96.9 % | 96.9 % |  | 99.7 % | 99.7 % |
| LT4 | 97.3 % | 97.3 % | 97.3 % | 99.7 % |  | 100 % |
| LT6 | 97.3 % | 97.3 % | 97.3 % | 99.7 % | 100 % |  |

### RNA-seq transcriptome genotyping and differential expression analysis

To further validate the genetic identity of the cybrids, we genotyped all cells using whole-genome SNP differences obtained from the RNA sequencing data (Supplementary Fig. 1A). While most cybrid genotypes matched their expected nuclear host cells, the LT4 (nucleus) x LE1 (mtDNA) heterocybrid clustered with the LE2 brown hare cell line. Due to the possibility of laboratory note confusion, this cell line was excluded from all subsequent analyses, leaving two reciprocal heterocybrid pairs and two homocybrid cell lines for further investigation. Overall, the cybrids had relatively minor changes in gene expression, with the cybrid cells clustering among the species represented by their nucleus (Fig. 1A).

**Fig. 1.**
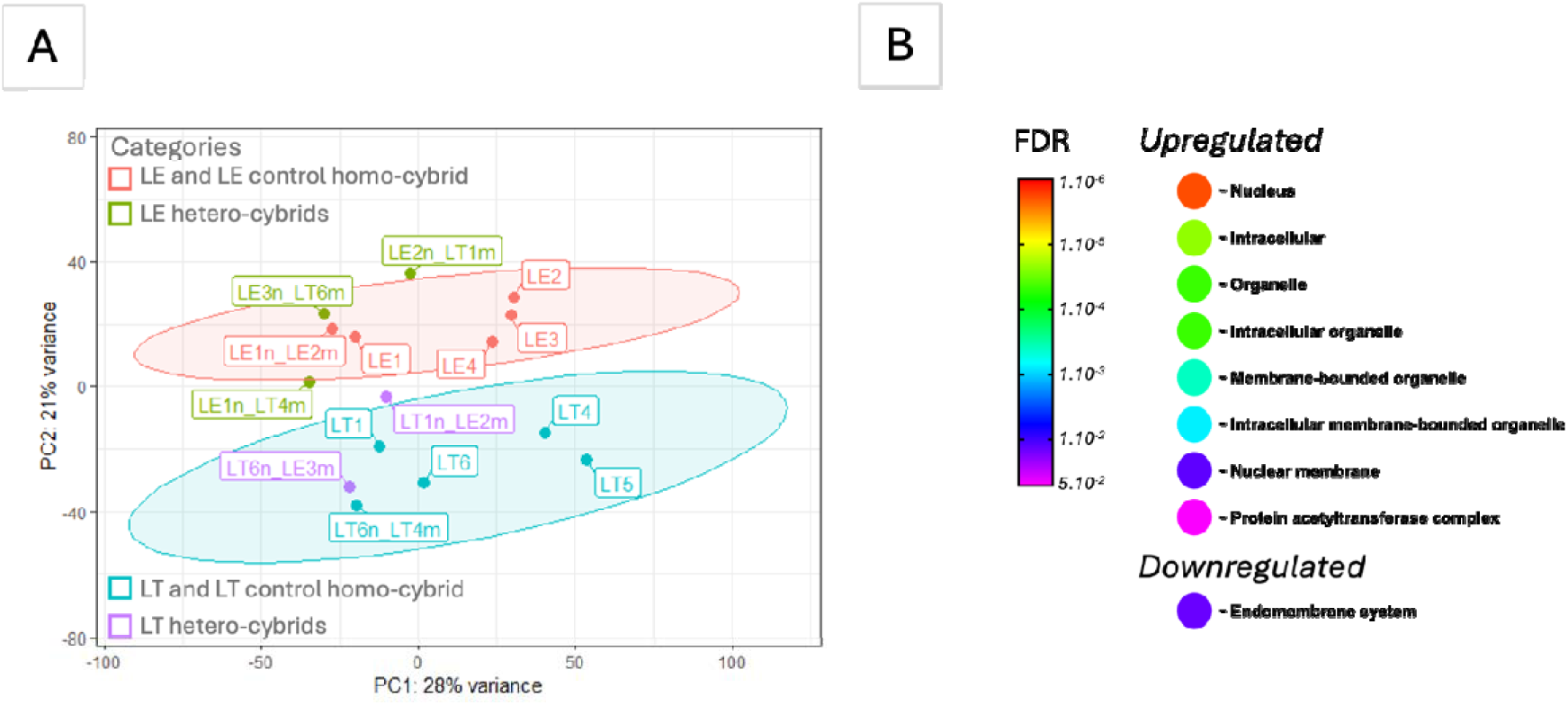
(A) Principal component analysis (PCA) of hare transcriptome revealed that cell lines cluster according to their nucleus identity. Two principal groups were identified with 95% CI: LE brown hare group (red cluster) and LT mountain hare group (blue cluster). n, nucleus; m, mtDNA. (B) STRING enrichment analysis. A significant subset of the transcripts upregulated in the xenocybrid cells are related to the nucleus and the nuclear membrane. Note that the “nuclear membrane” transcripts are also part of the “nucleus” category which are themselves included in the other upregulated categories presented in the figure. The transcripts downregulated in the xenocybrid cells show only weak association with the endomembrane system. See Supplementary Table 3 for additional information.

To investigate the global impact of nuclear-mitochondrial interchange on gene expression in the heterocybrids, we analysed the direction of transcriptional changes (increase or decrease) by comparing each heterocybrid to both parental cell lines. We then categorized each transcript into one of the 1024 possible combinations of up- and downregulation. For instance, transcripts that were upregulated in the heterocybrids compared to their nuclear parent but downregulated compared to their mitochondrial parent represented one such combination. Multinomial analysis followed by Bonferroni-corrected Chi2 tests revealed a significant deviation from random distribution for 9 categories, at all levels of gene expression thresholding (Supplementary Table 1). Genes systematically upregulated in the heterocybrids compared to both parental cell lines as well as the opposite category (genes systematically downregulated in heterocybrids) were dramatically overrepresented, indicating that the cybrid generation process could alter gene expression, possibly due to the selection reagents used (see methods “Gene expression change analysis”). STRING [57] functional protein association network analysis (Fig. 1B) of the genes upregulated in heterocybrids revealed strong enrichment in the nuclear compartment-associated genes. In contrast, the genes systematically downregulated in the heterocybrids were enriched in proteins associated with the endomembrane system.

Apart for the previously reported species differences [27], the comparative transcriptome analysis revealed genes with similar expression changes (up- or downregulated) across all cybrid types (both hetero- and homocybrids) relative to the parental cell lines, likely reflecting cellular alterations induced by the cybridization process or generic transcriptional responses to genome mixing (Supplementary Table 2, Supplementary Fig. 2). Most of the downregulated genes encoded ribosomal subunits and genes involved in protein synthesis, including translation initiation factors, as well as many transcription factors, particularly those coding for zinc finger and histone-like proteins. Interestingly, the expression of certain ubiquitin-conjugating enzymes and ubiquitin-protein ligases, which tag proteins targeted for degradation with ubiquitin, was also reduced. Overall, these results suggest a general repression of transcription, translation, and protein turnover in the cybrids. Moreover, in all cybrids genes involved in organization and dynamics of actin and microtubules were commonly downregulated, which could impact cell motility and genome integrity. Examples are ACTA1, an actin isoform, FES, a tyrosine kinase with a role in the regulation of the actin cytoskeleton, microtubule assembly, cell attachment and cell spreading, and DMTN (dermatin) having a role in modulating actin dynamics and formation of cell protrusions, such as filopodia and lamellipodia, necessary for cell spreading, motility and migration [58]. Likewise, CCDC8 and OBSL1, core components of the 3M complex required to regulate microtubule dynamics and genome integrity [58], were downregulated as well. Simultaneously, numerous core genes involved in cell adhesion (e.g. PRELP, ISLR, TNFRSF18, THBS4, ITGA11, VIT, SSPN), extracellular matrix production and organisation, such as collagens (e.g. COL4A5, FMOD), were highly downregulated, suggesting that adhesion properties in all cybrids might be reduced to facilitate cell motility. Among growth-associated genes, hepatocyte growth factor (HGF) and leptin (LEP) were commonly found downregulated. Additionally, GREM1, an antagonist of bone morphogenic protein (BMP) signalling [59], had reduced transcription levels in all cybrids compared to the original hare fibroblast cell lines. We were also interested to see if the exchange of mitochondria caused any changes in the mitochondrial compartment or cellular metabolism in general. We found upregulation of the mitochondrial 10 kDa heat shock protein (HSPE1 or Hsp10) with simultaneous downregulation of 60 kDa heat shock protein (HSPD1 or Hsp60). Together, these chaperonins facilitate correct folding of imported proteins into the mitochondrial matrix, promoting their refolding and proper assembly into polypeptides [60]. Likewise, the mitochondrial alpha subcomplex subunit 9 of Complex I (NDUFA9) and subunit 6 of Complex III (QCR6) were upregulated, while the β-subcomplex subunit 4 of Complex I (NDUFB4) and subunit 7 of Complex III (UQCRB) were downregulated. Two mitochondrial ATP synthase (Complex V) subunits, ATP5MK and ATP5MC3, together with some of the mitochondrial subunits of Complex IV (COX7B, COX5A, COX6A1, COX7A2, HIGD1A) were downregulated. Transcriptome results also showed inhibition of expression of mitochondrial contact site and cristae organising system (MICOS) complex subunit MIC10-like in all cybrid cells. The MICOS complex, located in the mitochondrial inner membrane, plays a crucial role in maintaining crista junctions, organizing inner membrane architecture, and forming contact sites with the outer membrane [61]. Similarly, TUSC2 involved in the maintenance of mitochondrial Ca^2+^ homeostasis and mitochondrial membrane potential (MMP)[62] was downregulated in all cybrids. Interestingly, considering that MMP is a driving force for Complex V for ATP production, also the levels of the mitochondrial SLC25A3 ion transporter were depleted in cybrid cells. This transporter provides phosphate ions essential for ATP production [63], as well as copper ions required for Complex IV assembly [64]. Furthermore, the levels of ARMCX2 shown to regulate mitochondrial dynamics in a number of cell types [65, 66] were similarly downregulated in all cybrids. Among metabolism-associated genes, we found downregulation of mitochondrial ACSS3 and ECHDC2 enzymes. The first one catalyses the synthesis of acetyl-CoA from short-chain fatty acids (FA) [67], while the second one enables enoyl-CoA hydratase activity and is predicted to be involved in FA β-oxidation [68].

Next, we identified 295 differentially expressed genes (DEGs) between LE heterocybrids (LE (nucleus) x LT (mtDNA)) and LE parents + LE control homocybrids (LE1 (nucleus) x LE2 (mtDNA), Fig. 2A, Supplementary Table 3). In LE heterocybrids, several pro-apoptotic factors, namely DAPK2, PRUNE2 and AIFM3, were downregulated, as well as GSDMD which promotes pore formation not only in the plasma membrane, but also in lysosomes and mitochondria [69]. Additionally, these cells showed decreased levels of NOX5, which generates superoxide from molecular oxygen utilizing NADPH as an electron donor [70]. We also observed reduction of NAPRT which encodes an enzyme that catalyses the biosynthesis of NAD from nicotinic acid and is associated with oxidative stress prevention [71]. The interaction between genes involved in cell proliferation and migration was multifaceted as well, involving downregulation of genes that regulate actin cytoskeleton, such as RFLNA, FBXL22 and WASF1. Several critical mediators of cell adhesion, namely protocadherin PCDH20, claudin CLDN7 and integrin ITGB3, showed decreased expression. Similarly, expression levels of myostatin (MSTN) that activates primary fibroblasts and stimulates collagen production [72] and dermatopontin (DPT) that facilitates adhesion of dermal fibroblasts [73] were strongly upregulated. Regarding genes associated with mitochondrial metabolism, we found a notable upregulation of MSS51, a mitochondrial translational activator required for assembly of respiratory chain Complex IV [74]. This finding was particularly interesting as it contrasted with the downregulation of several COX subunits, observed in all cybrid cell lines. In line with the OXPHOS downregulation in cybrids, several mtDNA encoded subunits of Complex V (ATP6, ATP8) and Complex I (ND3, ND6) had low expression levels in the LE heterocybrids. In these heterocybrids three genes involved in fatty acid metabolism were strongly downregulated as well, specifically a regulator of adipocyte differentiation PPARG [75] and ACAD10 involved in FA β-oxidation [76].

**Fig. 2.**
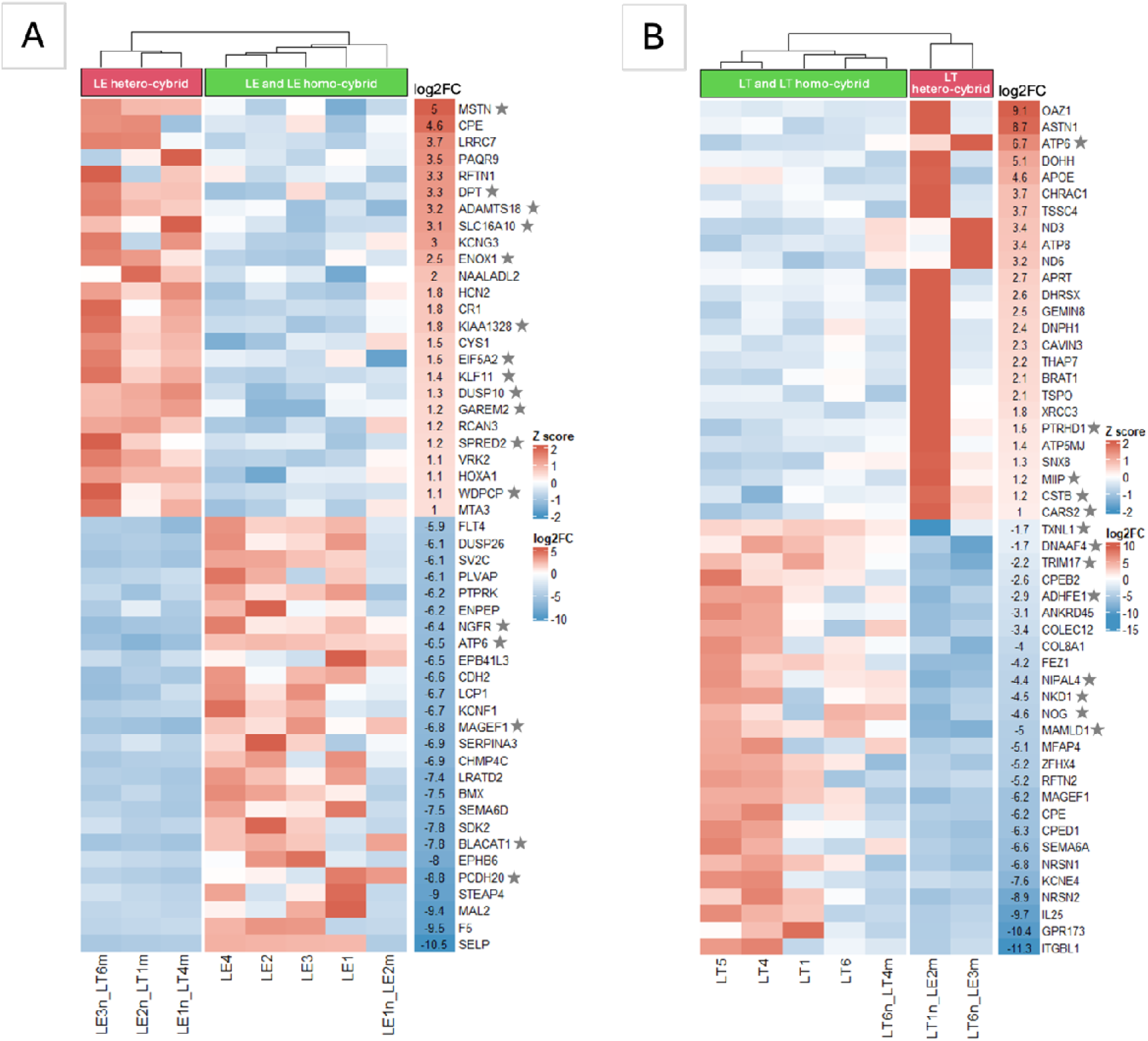
Heatmaps of top 25 upregulated and downregulated orthologs in (**A**) LE brown hare group and in (**B**) LT mountain hare group. Stars indicate “high confidence genes” with differential z-scores in heterocybrids compared to control homocybrids and respective parental cell lines. n, nucleus; m, mtDNA.

In total, 123 DEGs were identified between LT heterocybrids (LT (nucleus) x LE (mtDNA)) and LT parents + LT control homocybrid (LT6 (nucleus) x LT4 (mtDNA)), Fig. 2B, Supplementary Table 4). The LT heterocybrids showed inconsistent patterns of upregulated genes (Fig. 2B). For example, while the LT6 (nucleus) x LE3 (mtDNA) heterocybrid had elevated levels of mitochondrial encoded Complex V (ATP6, ATP8) and Complex I (ND3, ND6) subunits, the LT1 (nucleus) x LE2 (mtDNA) heterocybrid showed upregulation of ATP5MJ subunit of Complex V as well as cell proliferation and migration associated CHRAC1 [77] and OAZ1 [78]. In addition, these cells also showed upregulation of two DNA repair associated factors, BRAT1 and XRCC3, while the LT6 (nucleus) x LE3 (mtDNA) heterocybrid did not. Despite these differences, the two LT heterocybrids had also common features, such as the downregulation of the NKD1 inhibitor, involved in positive regulation of cell proliferation [79].

### Mitochondrial DNA copy number and OXPHOS protein levels

The downregulation of the mtDNA-encoded Complex I and Complex V OXPHOS subunits in the cybrids prompted us to investigate whether these cells had also an altered mtDNA content and mitochondrial protein expression. The parental brown hare fibroblasts (LE) exhibited a higher mtDNA copy number compared to those of the mountain hare (LT) (Fig. 3A). Cybrids with LE nucleus retained a similar mtDNA copy number to the parental LE cells, whereas a significant reduction in mtDNA copy number was observed in the LT heterocybrids, also when compared with the LT parental cells. As previously reported [27], the Complex I protein levels in mountain hares were higher than in brown hares, while the opposite was true for Complex IV (Supplementary Fig. 3).

**Fig. 3.**
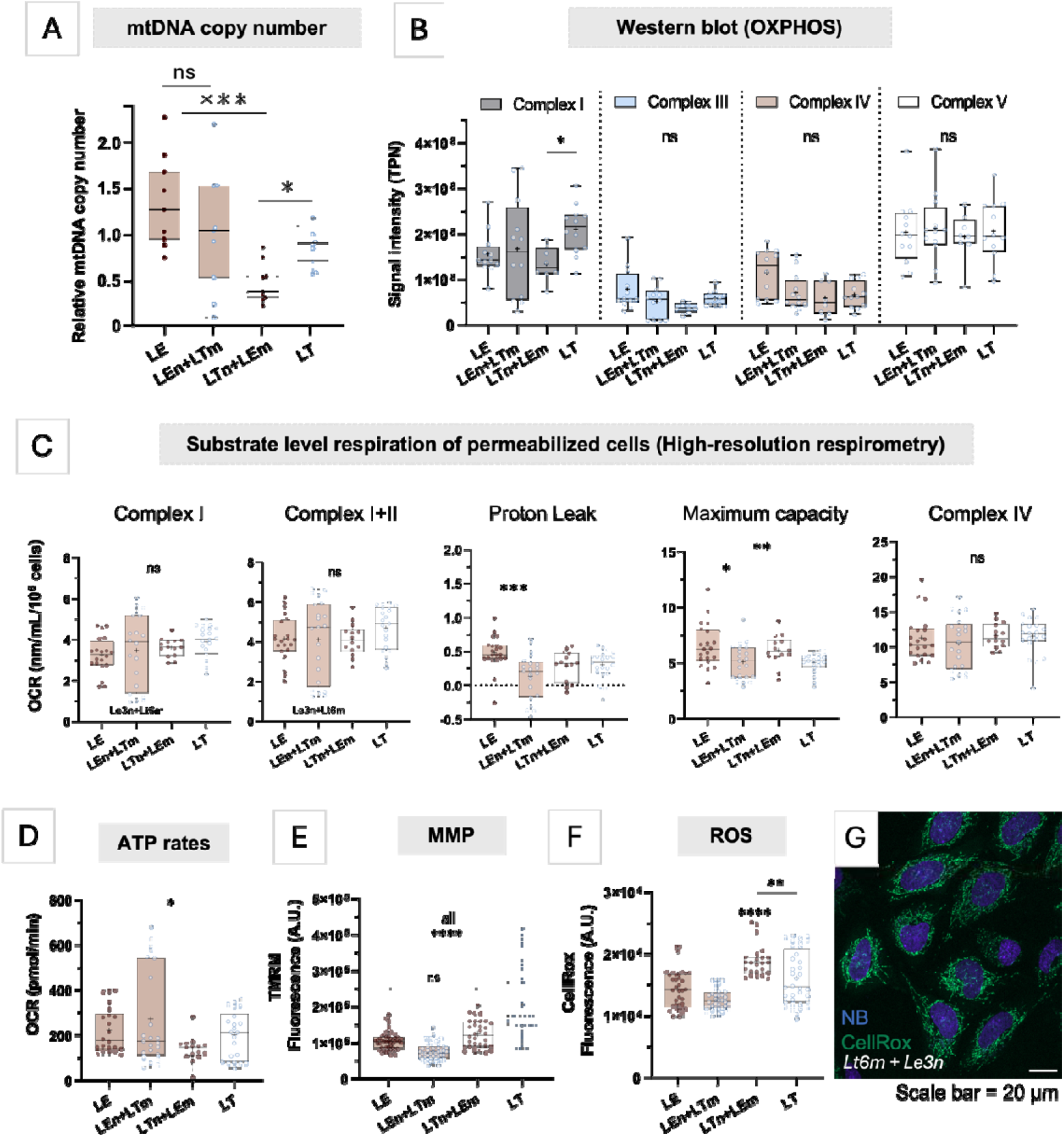
(A). Brown hare fibroblasts (LE) demonstrated higher mtDNA copy number compared to the mountain hares (LT). (B) OXPHOS protein expression measured with Western blot. n, nucleus; m, mtDNA. (C). Respiration parameters assessed in permeabilized cells using high resolution respirometry. OCR, oxygen consumption rate. (D) Total ATP production rates assessed with Seahorse Real-Time ATP Rate assay. (E) Mitochondrial membrane potential (MMP; LE vs LEn+Lem ns, all others *p* <0.0001) and (F) reactive oxygen species (ROS) production. (G) Representative confocal microscopy image of live cells stained with a ROS sensitive dye (CellRox) in green and a nuclear marker (NucBlue, NB) in blue. n, nucleus; m, mtDNA. Data presented as individual datapoints as well as boxplots with IQR, median (line) and mean (+) ± SD. *p*-value: * - <0.05; ** - <0.01; *** - <0.001; **** - <0.0001.

Despite differences in transcripts and mtDNA copy numbers, the protein levels of Complex V did not change significantly in any of the cybrids, as assessed by Western blot (Fig. 3B, Supplementary Fig. 3C, E, G, K). No significant differences were observed in the levels of VDAC1 (Supplementary Fig. 5A), a known regulator of mitochondrial apoptosis and metabolism [80].

In the heterocybrids, the Complex I protein levels showed a similar pattern of variation as the mtDNA copy number (Fig. 3B). In the LT6 (nucleus) x LE3 (mtDNA) heterocybrid, which previously demonstrated upregulation of ND3 and ND6 mitochondrially encoded subunits, the Complex I protein levels were not significantly altered (Supplementary Fig 3G), while the LT1 (nucleus) x LE2 (mtDNA) heterocybrid had decreased levels of Complex I compared to its LT1 parent (Supplementary Fig. 4E). The decreased mtDNA content combined with the lack of transcriptional upregulation may explain the observed decrease in Complex I levels in LT heterocybrids (Fig. 3B). The opposite situation was observed in all LE (nucleus) x LT (mtDNA) heterocybrids, which preserved mtDNA copy numbers, but showed decreased transcript levels of Complexes I, III and V. However, only in the LE3 (nucleus) x LT6 (mtDNA) heterocybrid were the protein levels of Complexes I and III strongly reduced (Supplementary Fig. 3G).

#### Oxidative metabolism in cybrid cells

Potential incompatibilities between the nuclear and mitochondrial compartments in species hybrids would be expected to manifest as OXPHOS dysfunction. In permeabilized cells, no major differences were found when comparing oxygen consumption in mitochondria between the heterocybrids and their parental cell lines (Fig. 3C, Supplementary Fig. 4). However, the respiratory chain capacity of the LE3 (nucleus) x LT6 (mtDNA) heterocybrid was greatly decreased compared to its parental cell lines and the reciprocal combination (Supplementary Fig. 5). Seahorse respirometry of intact cells confirmed this observation (Supplementary Fig. 4) and revealed a decreased basal respiration rate in both LE3 x LT6 heterocybrids (Supplementary Fig. 5 B). Additionally, the LE2 (nucleus) x LT1 (mtDNA) heterocybrid had strongly decreased maximal and spare respiratory chain capacity (Supplementary Fig. 5 C-D), although its basal respiration was within parental levels (Supplementary Fig. 5 B) and no negative alterations were found in permeabilized cell respiration (Supplementary Fig. 6).

In conclusion, respiration was lowered or unchanged in the cybrids compared to the parental cell lines, and the overall mitochondrial ATP production rate, despite showing considerable variation between cell lines, was not altered in cybrids (Fig. 3D), with the exception of the LE3 x LT6 heterocybrid (Supplementary Fig. 7). Pairwise comparisons showed non-systematic effects, underscoring the predominance of inter-individual variability.

### Mitochondrial membrane potential and ROS in cybrid cells

Respiratory chain activity is intimately linked to mitochondrial membrane potential (MMP) and reactive oxygen species (ROS) production [81]. MMP was lower in all cybrids compared to the nuclear parent, being most affected in the LT (nucleus) + LE (mtDNA) heterocybrids (Fig. 3E). This effect was found in all heterocybrids, but also in homocybrids (Supplementary Fig. 8A-D), suggesting MMP decrease is a generic consequence of cybrid generation. Notably, LE (nucleus) homo- and heterocybrids showed the highest MMP sensitivity to Complex I inhibition (Supplementary Fig. 8C). In contrast, ROS production was decreased or unchanged in LE (nucleus) + LT (mtDNA) heterocybrids as well as in homocybrids while significantly increased in LT (nucleus) + LE (mtDNA) heterocybrids (Fig. 3F). Pairwise comparisons between each cybrid and its parental cell lines confirmed these effects (Supplementary Fig. 8E).

### Metabolic alterations in cybrid cells

To examine metabolic preferences in the cybrids, we grew cells on substrate-coated microarrays, thereby testing their ability to metabolize each of the 185 carbon and nitrogen substrates. We observed no statistical differences in the substrate preference among studied cell lines; nor when comparing the cybrid and parental cell lines based on the nuclear or mitochondrial donor. Commonly preferred and unfavoured substrates were shared among all cell lines (Supplementary Fig. 9A, B), indicating a consistent metabolic preference in fibroblasts of both hare species. The most preferred carbon substrates listed in the order of preference were α-D-glucose, D-maltose, D-mannose, D-fructose-6-phosphate, inosine, D-fructose, α-keto-glutaric acid, maltotriose, D-salicilin, 3-O-methyl-D-glucose and uridine; while the most preferred nitrogen substrates were L-glutamine, L-arginine-glutamine and L-asparagine-glutamine.

Given the observed downregulation of several genes related with FA metabolism, especially in LE (nucleus) x LT (mtDNA) heterocybrids, we investigated the ability of these cells to utilize long chain fatty acids (LCFAs) for β-oxidation. However, only the LT6 (nucleus) x LE3 (mtDNA) heterocybrids showed increased capability of LCFA-driven respiration (Fig. 4A).

**Fig. 4.**
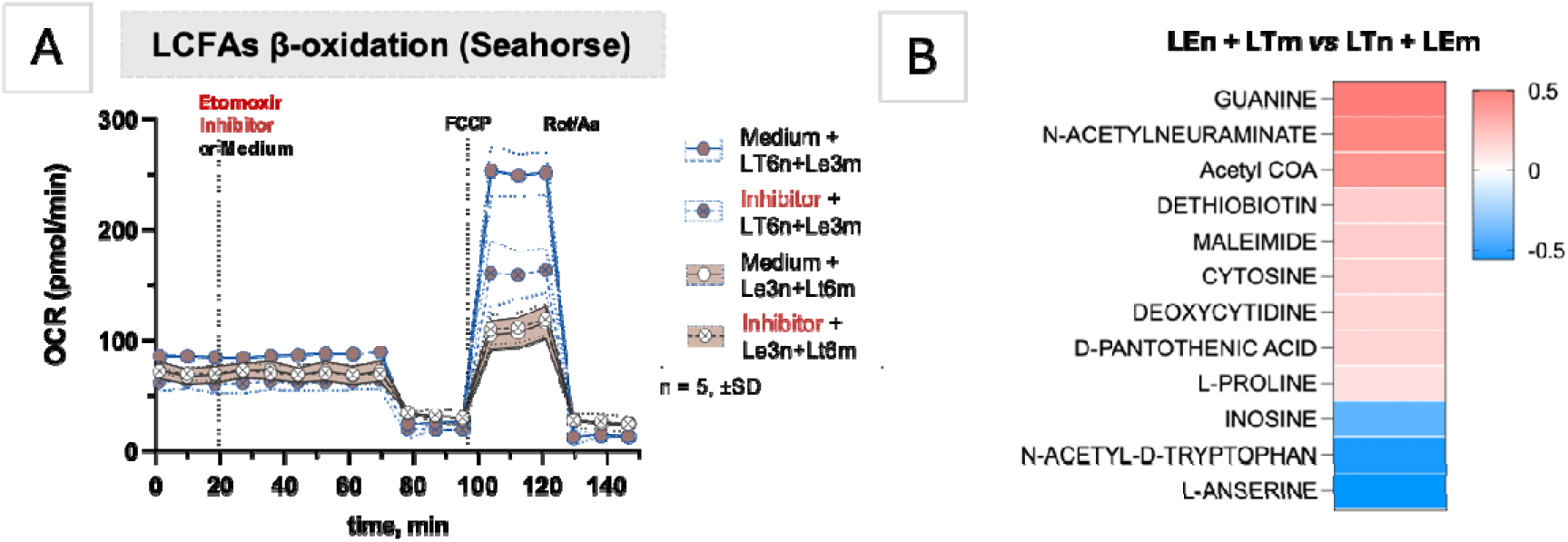
(A) Seahorse Long Chain Fatty Acids (LCFAs) Oxidation Stress assay revealed reliance on endogenous LCFAs for mitochondrial beta oxidation in LT6 (nucleus) x LE3 (mtDNA) heterocybrid, while the reciprocal heterocybrid did not demonstrate such a demand. (B) Of 130 identified metabolites with targeted metabolomics, 12 were found significantly different (FDR *p*-value < 0.05) between heterocybrids with LE nucleus/LT mitochondria and LT nucleus/LE mitochondria. n, nucleus; m, mtDNA.

To further investigate potential metabolic changes, targeted metabolomics was conducted and metabolic profiles of the reciprocal heterocybrids were compared. Overall, the heterocybrid lines with LE nuclei showed elevated levels of most measured metabolites (103 out of 130, Supplementary Table 5). In contrast to the transcriptome, heterocybrids belonging to the same parental cell lines clustered together rather than with the same species of their nuclear origin (Supplementary Fig. 9C). Amongst the metabolites present at significantly different levels based on the species of origin of the nucleus (Fig. 4B), anserine (FDR *p* = 0.04), n-acetyl-tryptophan (FDR *p* = 0.002), pantothenate (FDR *p* = 0.006), and acetyl-CoA (FDR *p* = 0.04) are all related to acetyl-CoA biosynthesis or to its regeneration through protein deacetylation and pantothenate pathway. Including LT and LE homocybrids in the analysis, respectively in the LT nucleus and LE nucleus groups, further increased the significance of the differences in the level of anserine (FDR *p* = 0.003), pantothenate (FDR *p* = 0.005), and acetyl-CoA (FDR *p* = 0.03), suggesting that differences in these metabolites are linked to the nuclear origin.

### Migratory and proliferative capacity of the cybrid cells

We confirmed our previously findings showing that brown hare fibroblasts exhibit greater wound healing capacity than the mountain hare ones [27]. In the cybrids, the wound closure rates correlated with the species of the nuclear donor (Fig. 5A). However, pairwise comparisons between reciprocal heterocybrids and their parental cell lines revealed strong individual differences (Supplementary Fig. 10A-B).

**Fig. 5.**
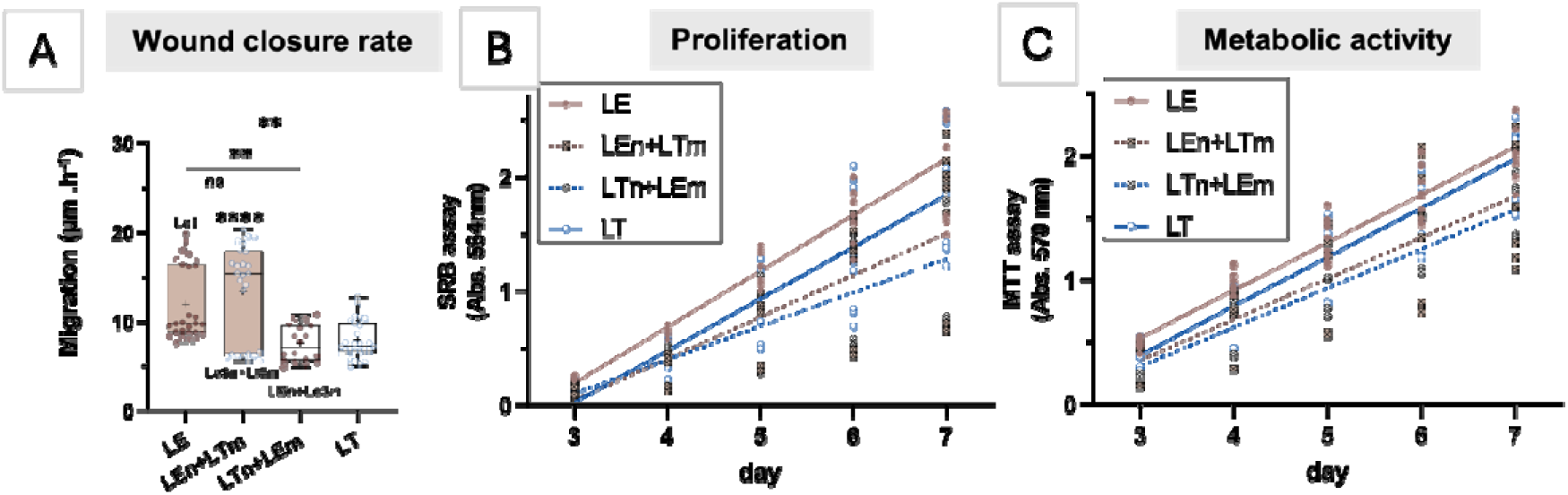
(A) Wound healing assay. (B) Cell proliferation measured with SRB assay and (C) metabolic activity measured with MTT assay. Each dot represents the measured cell lines at a particular time point connected with fitted linear regression curve. n, nucleus; m, mtDNA. Data presented as individual datapoints as well as boxplots with IQR, median (line) and mean (+) ± SD. *p*-value: * - <0.05; ** - <0.01; *** - <0.001; **** - <0.0001.

To measure cell growth, we employed the Sulforhodamine B (SRB) assay. SRB binds to cellular proteins, allowing us to estimate cell proliferation through repeated measurements taken at defined intervals from populations of cells seeded simultaneously at the same density [82]. Additionally, we applied the MTT assay that measures NADH as a proxy for cellular metabolic activity and thereby only detects viable cells. The MTT assay essentially led to the same observations as the SRB assays, although with different significance values (Fig. 5B, C). The proliferation of heterocybrids was generally decreased compared to the parental cell lines (*p* < 0.0001). However, we observed strong inter-individual differences (Supplementary Fig. 10C, D, Supplementary Table 6). Notably, the LE3 x LT6 reciprocal cybrid pair demonstrated decreased wound closure rates and proliferation compared to their parental cell lines (Supplementary Fig. 10). The LT control homocybrid had lowered proliferation rate, while the migration rate was similar compared to the nuclear donor cell line. In contrast, the wound closure rate of the LE homocybrid was elevated compared to both parents.

### Morphological features of the cybrid cells

Fluorescence imaging of the cells combined with cell morphometry analysis showed that mountain hare fibroblasts possess a larger cell size and larger nuclei than brown hares (Fig. 6A-C, Supplementary Fig. 11). This interspecific difference in nucleus size is intriguing, considering that the brown hare genome is slightly larger (2.9 *vs* 2.7 Gb) [49, 83]. Despite the nuclear size difference, there were no differences in the shape of the nucleus (Fig. 6D). In the cybrid cells, cellular size was similar to the nuclear parent, or even slightly increased (Fig. 6B). However, when comparing each cybrid to its specific parental cell lines, these influences appeared weak and non-systematic (Supplementary Figs. 12–14).

**Fig. 6.**
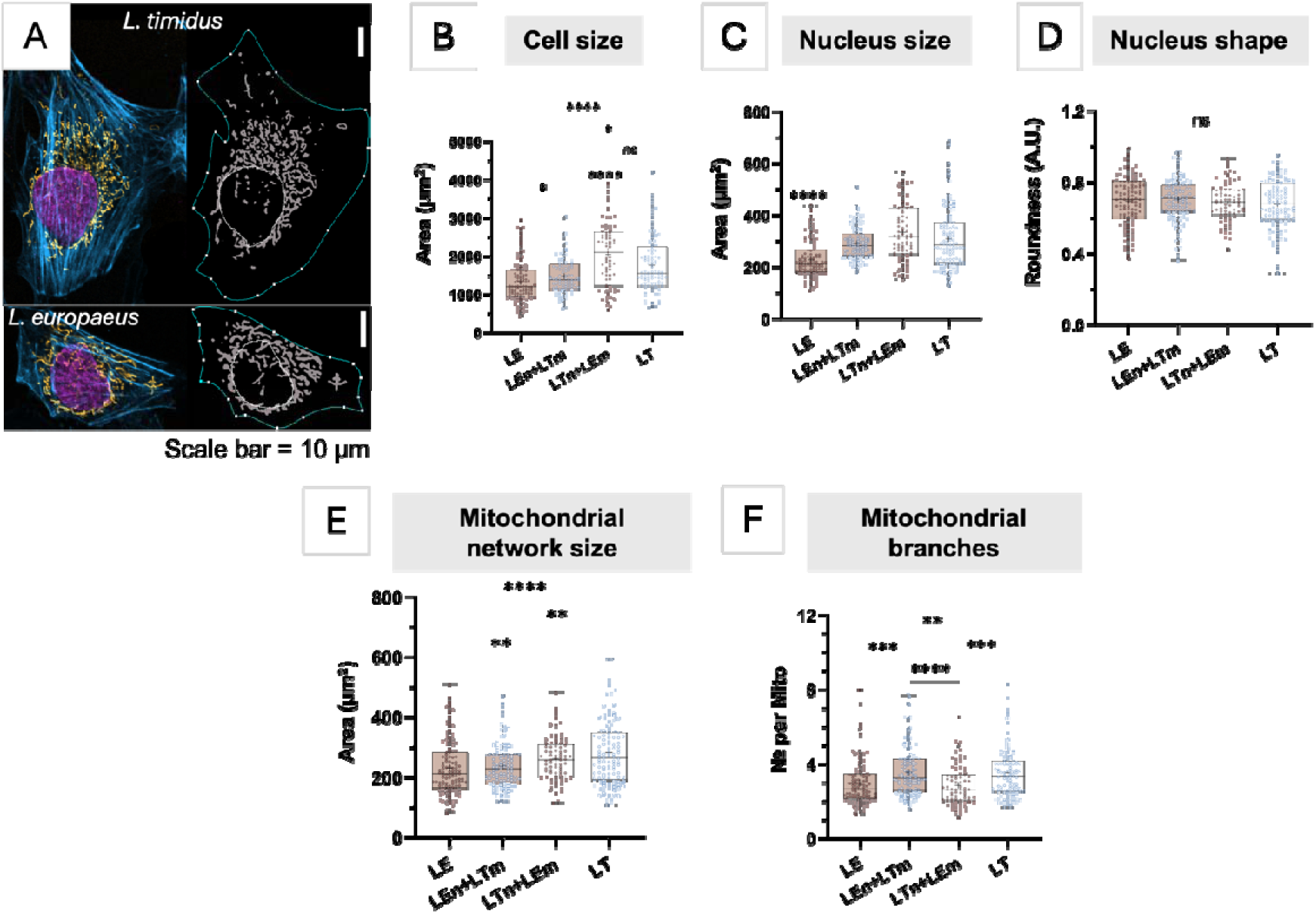
(A) Representative image of LT and LE fibroblast cells and an example of applied masks used to measure cell morphology parameters. (B-F) Measured cell morphology parameters. n, nucleus; m, mtDNA.

Suspecting that mitochondria-nucleus incompatibility could be more evident in its consequences on mitochondrial morphology, we focused our analysis on this aspect. Quantification of stained mitochondrial networks revealed that mountain hare fibroblasts possess larger and more branched mitochondrial networks compared to brown hare cells (Fig. 6E–F, Supplementary Fig. 11). In heterocybrids, mitochondrial network size resembled that of their nuclear parent, while mitochondrial connectivity was more similar to the mitochondrial donor (Supplementary Fig. 12). However, as observed in other comparisons, pairwise analysis of parent-cybrid variations revealed substantial variability between cybrid cell lines (Supplementary Fig. 13 and 14).

### Correlation of observed phenotypic features in cybrid cells

To further understand the connections between ROS, MMP and cell biology, we performed a comprehensive correlation analysis of all measured variables (Fig. 7). Expectedly, permeabilized cell respiration through CI, CI+II and CIV correlated positively with each other and with CI protein levels as assessed by WB. Also, the two assays (SRB and MTT) designed to measure cell growth correlated positively with each other. ROS levels were negatively correlated with cellular NADH content as measured by the MTT assay and ATP-coupled respiration in intact cells, all general indicators of poor cellular fitness. Mitochondrial membrane potential was positively correlated with the size of the nucleus.

**Fig. 7.**
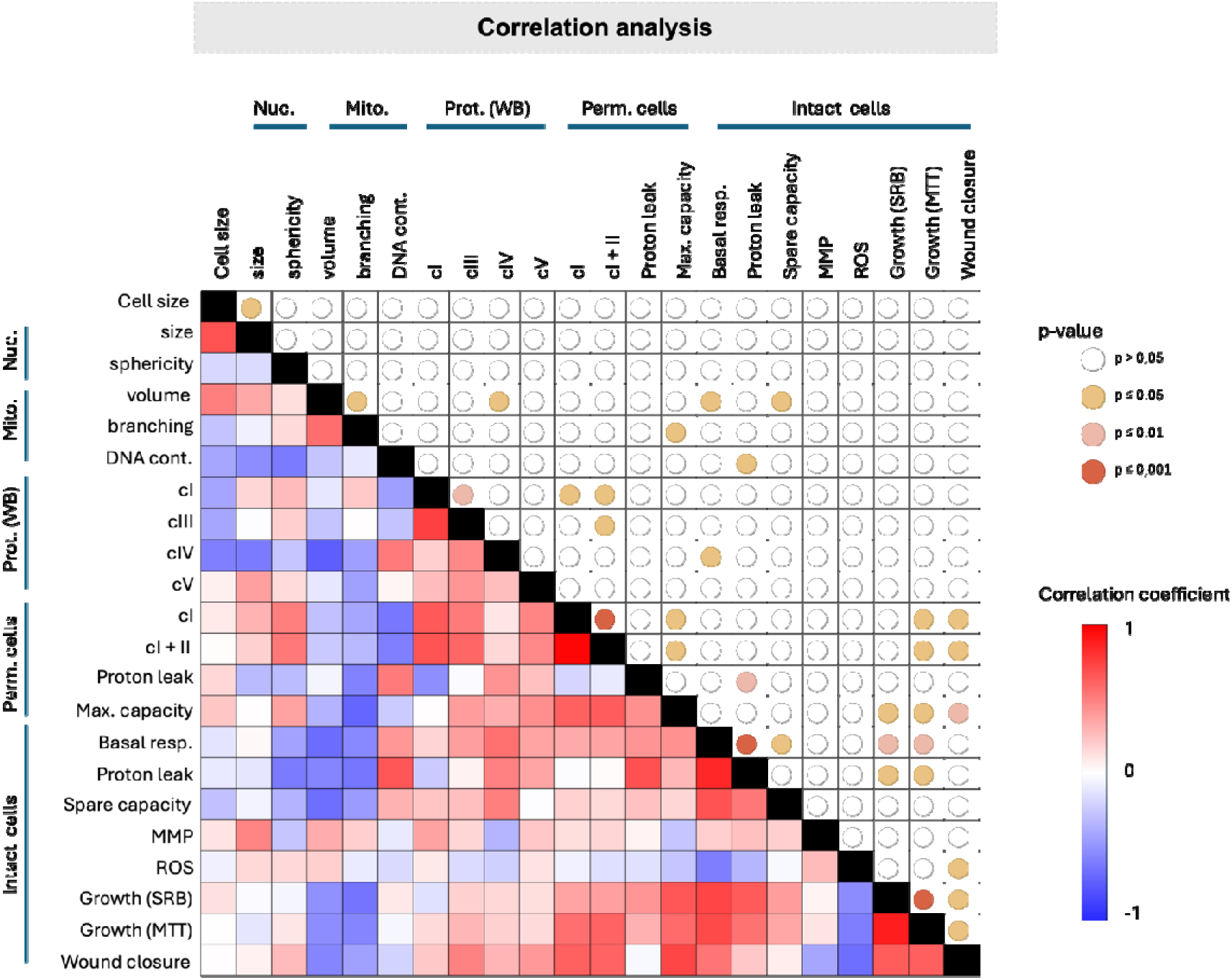
Correlation matrix showing potential interrelations between measurements in Fig. 1-6. The left and lower cells display the Pearson correlation coefficients for each comparison, with blue indicating negative and red indicating positive correlations, as detailed in the “correlation coefficient” legend on the right. The upper and right cells display uncorrected two-tailed p-values for the same comparisons, with significance levels indicated in the “p-value” legend at the top right.

## Discussion

The technology to generate cytoplasmic hybrid cells (cybrids), where a ρ0 host cell line is repopulated with mitochondria and mtDNA from different donor cells, was established already in the 1980s [32]. Cybrid cells have been used extensively to study the effects of different mtDNA haplotypes on cellular respiratory competence, mostly utilizing patient mutant mtDNAs to understand the pathological mechanisms behind mitochondrial diseases [84–89]. Additionally, interspecies xenomitochondrial cybrids have been studied, focusing on human-primate and rodent-rodent incompatibilities of the OXPHOS complexes [30, 90, 91]. Generally, interspecies cybrids have lower respiration rates, with Complex I being most affected and the degree of impact correlating with increasing evolutionary distance between species [30, 90]. When the evolutionary distance becomes too large, such as between humans and orangutans, the mtDNA-encoded OXPHOS subunits are no longer compatible with the nuclear DNA and fail to sustain functional mitochondrial respiration.

In this study, we used interspecific cybrids between mountain hare (LT) and brown hare (LE) to investigate the effects of introgressed mtDNA and its potential role in reinforcing species barriers among hares. We hypothesized that species incompatibilities between the mitochondrial and nuclear genomes would be evident not only in OXPHOS function, but also in other aspects of cellular fitness, potentially revealing mechanisms of hybrid breakdown in hares. Although simplified, our model aimed for increased realism by using cell lines from three different individuals per species, representing wild animals with greater genetic diversity than standard inbred model organisms, thus enhancing the relevance of our findings to natural hybridization scenarios. We also leveraged genetic differences between species and individuals to validate the phenotyping results. This approach proved valuable: despite using nuclear sex markers and mtDNA haplotypes to confirm cybrid cell identities during their generation (Table 1), genome-wide SNP genotyping from RNA sequencing later revealed that one LT (nucleus) x LE (mitochondria) cybrid might have been misidentified (Supplementary Fig. 1) and was therefore excluded from the analyses. It should be noted that similar genome-wide characterization of cybrids is usually not conducted, further underscoring the importance of genotypic verification when interpreting the results of phenotyping assays. Besides labelling mistakes of the parental cell lines, it is important to exclude that some nuclei from the mitochondrial donor cell survive the Actinomycin D treatment and outcompete true cybrids during subsequent passages in cell culture.

### Heterocybrids as mtDNA introgression models

Much interest has been focused on the adaptive significance of mtDNA haplotypes in humans and other animals, with studies exploring correlations between haplotypes and phenotypes related to longevity [92], endurance [93, 94], climatic adaptation [95], and disease susceptibility [96]. A recent study found that mitochondrial haplotype and tissue type significantly influence the somatic evolution of the mitochondrial genome as organisms age [97], suggesting that evolutionary processes within the organism can act to maintain compatibility between the mitochondrial and nuclear genomes. In this respect, our transcriptome, cell proliferation, respiration, measurements of ROS production, metabolic substrate assays and targeted metabolomics provide interesting insights into the global effects of altered mitonuclear interactions.

First, our transcriptome analysis showed that cybrid cells have a distinct transcriptome fingerprint, distinguishable from the differences observed between the two parental species (Figs. 1–2). In this respect, it is interesting that all cybrids shared increased expression of a subset of genes related to nuclear structure and function (Supplementary Table 1). This indicates that cybrid formation has previously undocumented, reproducible, long-term consequences maintained over many cell generations, a fact that should be accounted for when using cybrids as disease models or treatment [98]. It is plausible that the specificity of newly established mitonuclear interactions could shape differential patterns of gene expression. Changes in cellular metabolism can cause cross-generational alterations in gene expression *via* epigenetic modifications [99]. In fact, it appears that mitochondrial DNA haplotypes can influence DNA methylation patterns and thereby modulate chromosomal gene expression [100]. While the observed gene expression differences could reflect generic and biologically meaningful changes in cellular retrograde signalling due to altered bioenergetics, they might also result from the harsh chemical treatments required to generate cybrids. For instance, treatment with 2’3’-dideoxycytidine (ddC), an inhibitor of the mitochondrial DNA polymerase γ used to generate mtDNA-free ρ0 cells [101], can have adverse effects on cells [102, 103], which might cause permanent epigenetic changes. Similarly, the mitochondrial donor nuclei are destroyed with Actinomycin D, whose carryover in small doses could result in DNA damage also in the recipient cells and have chronic effects on gene expression, such as the activation of DNA damage responses as seen in LT1 (nucleus) x LE2 (mtDNA) (Supplementary Table 5). Importantly, the observations from un-pooled transcriptome data should be interpreted with caution, as replicate analyses of each specific sample were not conducted due to resource limitations.

### Fitness effects of heterocybrid condition on energy metabolism

The fact that introgressed mtDNA can persist in natural populations of hares long after the donor species has gone extinct [20] indicates that there are no significant incompatibility issues, at least between some species. This might not be surprising, given the relatively small differences between mountain hare and brown hare mtDNA coding sequences (Table 2), which account for a maximum of 59 amino acid differences between the species. As a comparison, the polypeptide coding genes of the human and the common chimpanzee mtDNA differ by 159 amino acid residues, contributing to a 41 % lower Complex I activity in human-chimp cybrids [90]. However, there is typically asymmetry in the introgression patterns. For example, brown hare mtDNA is extremely rare in wild mountain hare populations [16, 17]. While this can be explained by demographic or behavioural factors [19, 20, 104], selective pressure against introgressed brown hare mtDNA in mountain hares is also possible, especially as first-generation hybrids are known to occur [18, 21]. An interesting phenomenon to consider is the so-called “Mother’s curse”, where a selfish mtDNA haplotype has a deleterious effect in males but not in females due to its maternal inheritance [105]. It is noteworthy that a specific mtDNA haplotype has been associated with male infertility in captive brown hares [106], i.e. caused within the same species by a differing haplotype lineage. This raises the possibility that a mild mitochondrial defect in cells with LE mtDNA in an LT nuclear background might contribute to male hybrid sterility, particularly given that spermatogenesis and sperm motility are very sensitive to disturbances in OXPHOS function [107, 108]. A hybrid-specific Mother’s curse, potentially mitigated by increased nuclear gene complementation from the more compatible parental species, would be expected to manifest as reduced Y-chromosomal and X-chromosomal introgression compared to autosomes [109, 110]. However, this has not yet been shown for hares and warrants further investigation.

Our study suggests asymmetric incompatibility, as e.g. mtDNA copy number was clearly decreased and ROS production increased in LT(n)+LE(m), but not in reciprocal cybrids (Fig. 3A, F). Defects in mtDNA maintenance can be a consequence of mitochondria-nucleus incompatibility and could be mediated by the differences in the various elements of the non-coding region (NCR), which controls mtDNA replication and transcription [111]. Asymmetric incompatibility between nucleus-encoded regulatory factors of one species and control elements on the mtDNA of another species, but not *vice versa*, might explain the differences observed between reciprocal cybrids. It is noteworthy that while long repeat (LR) elements on the mitochondrial genomes of the two species show individual variation in repeat number, mountain hares maintain on average shorter repeat arrays than brown hares [31]. Furthermore, while the typical mountain hare mtDNA array length was maintained in LE (n) cybrids, the arrays showed instability in reciprocal cybrids, with new LR-length haplotypes appearing in heteroplasmy with the original [31]. While the LRs lack any obvious regulatory function, it is possible that their expansion or reduction is dependent on processes controlled by nuclear gene products, such as mtDNA replication. While such traits likely do not have any fitness effects, the existence of species-specific variation demonstrates that functionally meaningful sequence specificity between nucleus-encoded factors and mtDNA regulatory elements also exists.

The same feature might explain also the observed differences in mtDNA copy number. Although LT cells had a lower mtDNA copy number, this did not manifest in lower protein levels of the different complexes (Fig. 3B). This might not be surprising, as gene expression is not directly dependent on mtDNA copy number, and even dramatically low levels of mtDNA can maintain normal OXPHOS function [112]. However, the LTn+LEm cybrids with the lowest copy number had slightly reduced Complex I protein levels (Fig. 3B). We consider that this reduction is unlikely to be caused by the mtDNA copy number. As Complex I has the highest number of mtDNA-encoded subunits (ND1, ND2, ND3, ND4, ND4L, ND5 and ND6), which also exhibit the greatest amino acid divergences between the two species (Table 2), the observed reduction likely relates to stability issues within Complex I. Furthermore, Complex I is structurally and functionally the most dynamic part of the OXPHOS, regulating and maintaining metabolic homeostasis, as well as initiating stress responses, including ROS signalling under hypoxia [113]. Although cellular respiration and ATP production were not affected in the LTn+LEm cybrids (Fig. 3C,D), these cells showed lowered membrane potential and elevated ROS production (Fig. 3E–F), which might be attributable to the proton pumping functions and ROS-producing tendency of Complex I [113]. Curiously, the reciprocal LEn+LTm cybrids showed a lower proton leak and maximum respiratory capacity than the parental LE cells. A more detailed pairwise analysis of cell respiration both by Seahorse and high-resolution instruments (Supplementary Fig. 4), revealed that the performance of cybrids is usually an intermediate of the parental cells. This clearly demonstrates that both mitochondrial and nuclear components affect cell respiration. Our interpretation is that the potential incompatibility in the OXPHOS subunits does not affect the cell respiration or ATP production per se, but rather causes a more subtle functional defect that results – in some cybrid combinations – in decreased membrane potential and elevated ROS (Fig. 3E, F).

The greater the architectural complexity of multi-protein complexes [114], such as Complex I – which is encoded by both the mitochondrial and the nuclear genomes – and the more numerous the regulatory factors required for the assembling protein subunits [115] into functional entities, the higher the risk of failure due to subunit incompatibilities might be. In fact, recently available data strongly supports the Dobzhansky–Muller hypothesis about deleterious effects of specific genetic interactions in Complex I that may result in hybrid lethality [10]. Additionally to Complex I, also Complexes III and IV contribute significantly to proton (H) pumping and the regulation of mitochondrial membrane potential, which ultimately drives respiration and ATP production. In mammals, these complexes frequently interact dynamically [116], forming major respirasomes (supercomplexes) that facilitate electron transfer from Complex I via coenzyme Q to Complex III, and from Complex III via cytochrome c to Complex IV. The stability of these complexes can be compromised by even a single mutation in the *Cytochrome b* gene of Complex III [117]. The minor coding difference (∼1%) in *Cytochrome b* between the mountain hare and the brown hare could thus contribute to supercomplex instability, causing disruptions in electron transport and elevated ROS [118]. This possibility needs to be further tested in our cybrid model.

Apart from Complex I, most of the coding variation between mountain and brown hare were detected in Complex V. However, we did not observe differences in protein levels of Complex V in our cybrid models (Fig. 3B). Considering that Complex V is essential for ATP production and heat generation [119], its misassembly might be lethal and hence not observed in viable hybrids. On the other hand, as Complex V typically forms oligomeric chains rather than physically interacts with other OXPHOS complexes [120], its structural variation may be better tolerated, while potentially contributing to the observed differences in oxygen consumption (Fig. 4D) and possibly heat generation.

#### Metabolic shifts in energy metabolism and wound healing of cybrids with LE nucleus

Among metabolites showing significant differences depending on the nuclear species of origin, anserine, N-acetyl-tryptophan, pantothenate, and acetyl-CoA are all associated with the biosynthesis or regeneration of acetyl-CoA that occurs either through protein deacetylation or via the pantothenate pathway. Both acetyl-CoA and pantothenate levels were notably higher in LE (nucleus) x LT (mtDNA) heterocybrids (Fig. 4B). Acetyl-CoA, a key metabolite in cellular metabolism and bioenergetics, plays a central role across various biochemical pathways [121]. Meanwhile, pantothenate, also known as vitamin B5, is known to enhance wound healing and stimulate skin cell proliferation [122], potentially explaining the enhanced wound healing capacity observed in cell lines containing a nucleus of LE origin. Collectively, these findings – although preliminary – indicate increased metabolic activity in the LE (nucleus) x LT (mtDNA) heterocybrids.

As another interesting observation, the reciprocal hare cybrids were metabolically similar (Fig. S9), despite carrying nuclear and mitochondrial genomes from different species. This similarity may arise if metabolites transferred during the cybrid creation trigger epigenetic changes in the recipient cell, leading to metabolic reprogramming in the cybrids [123]. This effect contrasts with the metabolic priming and adaptive changes expected to occur in true species hybrids, where tissue-specific cellular metabolism develops during normal embryonic growth. In fact, studies on xenomitochondrial mice with relatively mild mitonuclear incompatibility have shown only modest changes in cellular physiology and gene expression, none of which were mitochondria-associated [91]. It should be noted that treatment-induced effects are likely unavoidable in cell culture, emphasizing the need for caution when interpreting cybrid data.

### Fitness effects of heterocybrid condition on cell mobility and structure

The wound closure rate is a measure of cell proliferation and migration, which both play important roles not only in wound healing, but also in tissue morphogenesis and cancer metastasis [124]. A surprising feature of our cybrids was that while they showed consistently lower proliferation rate and metabolic activity, their wound closure rate was significantly increased in the LE nucleus + LT mtDNA cybrids compared to the parental cells (Fig, 5). As cell motility or migration is dependent on the complex interplay between cytoskeletal dynamics, cell adhesion and signalling responses to chemotactic cues [125], it is challenging to speculate on the mechanisms underlying this enhanced motility in the cybrids.

The fact that mountain hare and brown hare fibroblasts differ in their morphology (Fig. 6) allowed us to monitor the effects not only of the mtDNA haplotype but also the cybrid generation process on cell shape, size and mitochondrial network structure. Despite some differences, cybrid cells retained the original morphology of their nuclear donor regardless of the mitochondrial donor identity. This is perhaps unsurprising, considering that all factors known to regulate the cell size and morphology are encoded by the nucleus. Additionally, the mitochondrial networks in cybrid cells did not differ significantly from the parental nuclear donors. While some cell lines showed increased mitochondrial branching (LE3n+LT6m, LT4n+LE1m), others had decreased (LT1n+LE2m) branching compared to the mitochondrial donor cells. Both mitochondrial hyperfusion and fragmentation have been extensively studied in the context of various pathological conditions reflecting deregulation of proteins responsible for mitochondrial fission–fusion dynamics [6]. Importantly, certain stressors, including Actinomycin D, can induce mitochondrial hyperfusion via L OPA1, MFN1 and SLP-2, which are supposed to stimulate mitochondrial ATP production and promote a pro□survival response [126]. However, while LE3n+LT6m and LT4n+LE1m cybrids showed increased mitochondrial network connectivity, their ATP production rates were decreased (Supplementary Figs. S1, S5).

## Conclusions

In the presented study, we have explored the impact of mitochondrial and nuclear genome interactions on cellular function by generating interspecific cybrids between mountain hares and brown hares. Through a combination of transcriptomics, metabolic assays, and functional assessments, we aimed to uncover how mtDNA introgression influences cellular fitness and compatibility in hybrids, thereby providing insights into mitonuclear interactions at species boundaries. Although LTn+LEm cybrids displayed reduced mtDNA levels, impaired complex I activity, and elevated ROS production - consistent with prior findings on interspecies cybrids - our results were generally mixed, with indications of potential metabolic carryover effects. These findings underscore the need for careful interpretation, as changes induced by cybrid generation may obscure the subtle effects caused by altered mitonuclear crosstalk. While not ideal for studying genetic compatibility in species hybrids, the phenotypic variation in cybrids provides valuable opportunities to explore the interactions between gene expression regulation and metabolic reprogramming, along with ROS production, membrane potential, stress responses and cell cycle regulation.

## Acknowledgements

The authors acknowledge Biocenter Finland and the Tampere Imaging Facility, Tampere University Flow Cytometry Facility and Cellular Respiration and Energetic Metabolism Facility, Tampere Finland, as well as Anni I. Nieminen at the Biocenter Finland and HiLIFE-funded FIMM Metabolomics Unit, Helsinki, Finland. Ms Anita Kervinen (UEF) is thanked for her valuable laboratory assistance.

## Author Contributions

JP, ED and SS contributed to the study conception and design. Material preparation, data collection and analysis were performed by KG, RT, SG, SS, ZF and ED. The first draft of the manuscript was written by JP and all authors commented on previous versions of the manuscript. All authors read and approved the final manuscript.

## Data Availability

The datasets generated for and analysed during the current study are available in the NCBI SRA repository, under the BioProject accessions PRJNA826339 and PRJNA1188074, for the parental and cybrid cell lines, respectively.

