## Supplementary information file for "Exploring mitonuclear interactions in the regulation of cell physiology: insights from interspecies cybrids"

**Supplementary Table 1. Comparison of gene expression between cybrids and their parental lines.** This table identifies groups of genes with consistent patterns of expression change across cybrids relative to their parental lines, significantly overrepresented compared to a random distribution. **Sheet 1:** Of 1,024 ($2^{10}$) possible gene-expression groups, nine (Groups A–F) are significantly overrepresented across all threshold levels. Columns B–K indicate the direction of gene-expression change for each comparison (+1: increased expression in cybrids; -1: decreased expression). Columns L–T provide statistical data (e.g., number of genes in each group, total genes tested, and significance) for thresholds of 0, 50%, and 2-fold changes. Columns U–Z present Gene Ontology (GO) enrichment analyses for each group in the absence of thresholding, with gene names provided for Groups A and B. **Sheet 2:** Lists genes in the nine significantly overrepresented groups without thresholding, including gene IDs and full names.

**Supplementary Table 2.** A list of differentially expressed confounding genes between cybrids and parental cell lines.

**Supplementary Table 3.** A list of differentially expressed high confidence genes in LE (nucleus) x LT (mtDNA) hetero-cybrids.

**Supplementary Table 4.** A list of differentially expressed high confidence genes in LT (nucleus) x LE (mtDNA) hetero-cybrids.

**Supplementary Table 5.** Differences in 130 metabolites identified through targeted metabolomics between LEnLTm and LTnLEm heterocybrids.

**Supplementary Table 6.** Cell line to cell line comparison of MTT and SRB assays.

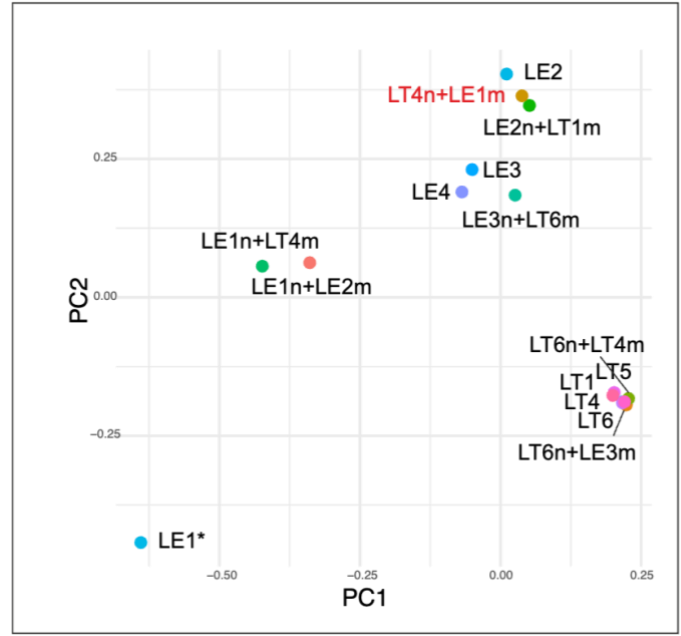

**Supplementary Fig. 1.** Validation of the cybrid cells by SNP genotyping of their transcripts. LT4n+LE1m cybrid (red) matched with LE2 parental cell line while all other cybrids clustered with their expected nuclear hosts. \*The parental LE1 is an outlier due to issues with SNP calls, however both cybrids with LE1 nucleus cluster together and with other LE cells. n, nucleus; m, mtDNA.

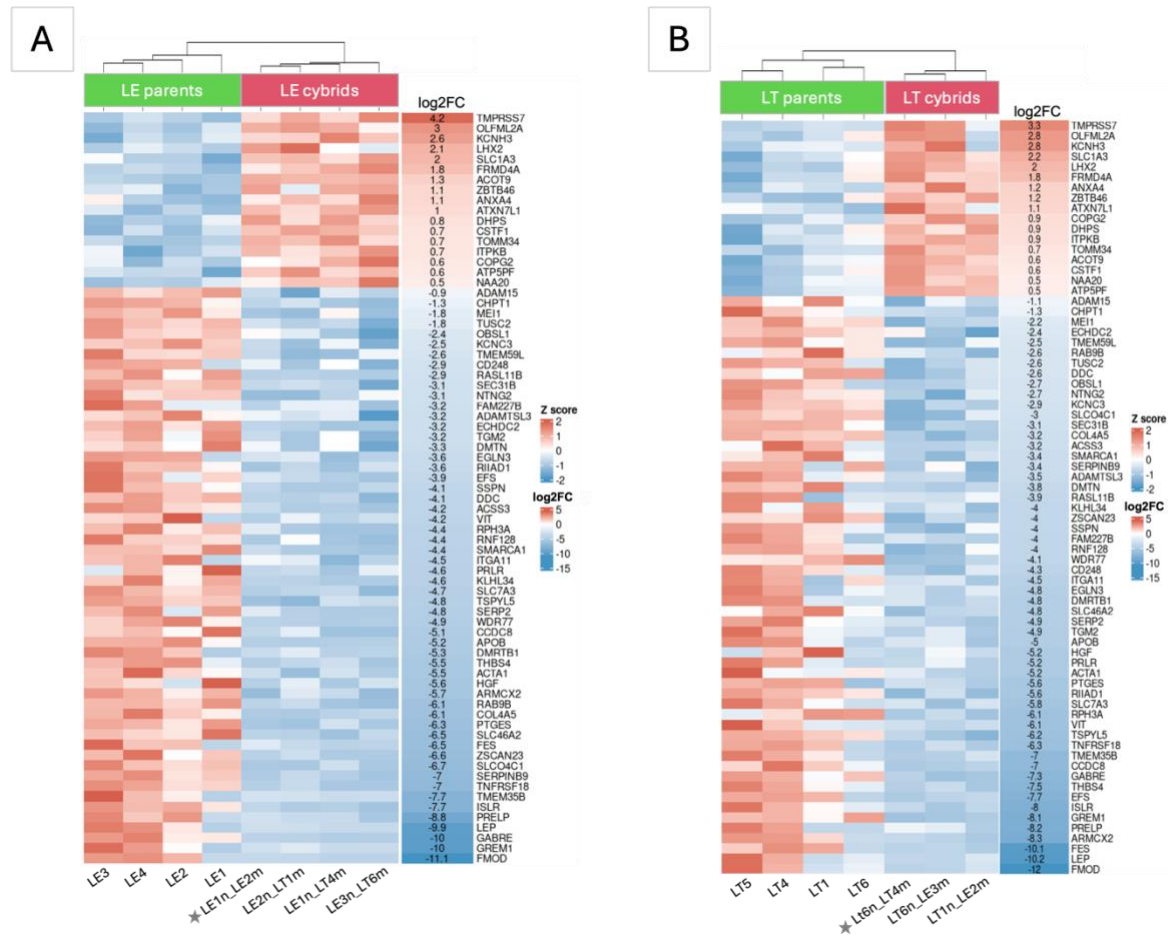

**Supplementary Fig. 2.** Heatmaps of all commonly changed orthologs (17 upregulated and 57 downregulated) in cybrids compared to parental cell lines (A) LE brown hare group, (B) LT mountain hare group. Stars indicate control homocybrids. n, nucleus; m, mtDNA.

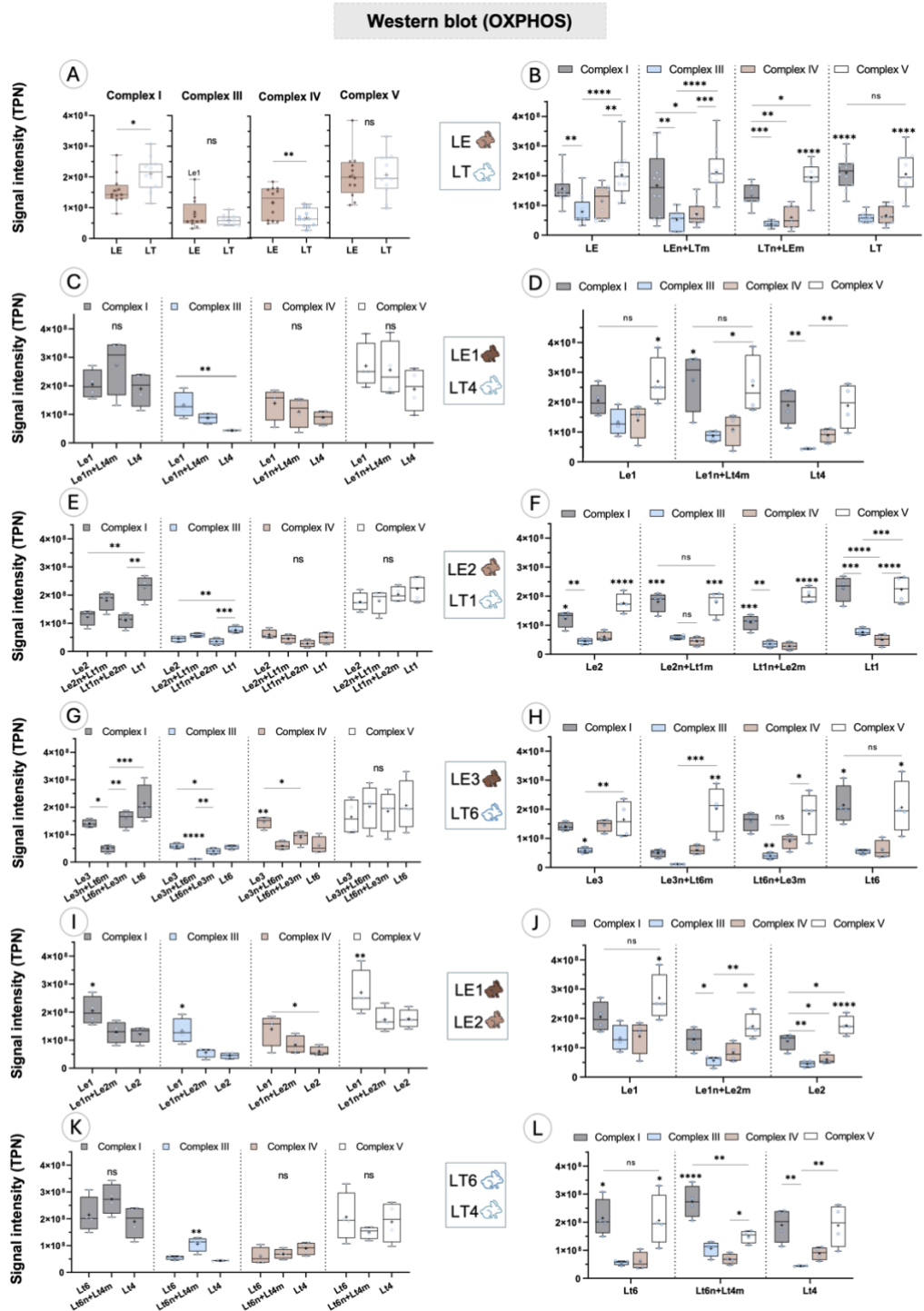

**Supplementary Fig. 3.** Western blot of OXPHOS proteins. **A, C, E, G, I, K** comparisons of Complex I, III, IV, V protein expressions between different cell lines. **B, D, F, H, J, L** comparisons of relative Complex I, III, IV, V protein expressions within the same cell line. n, nucleus; m, mtDNA. Data presented as individual datapoints as well as boxplots with IQR, median (line) and mean (+) ± SD. *p*-value: \* - <0.05; \*\* - <0.01; \*\*\* - <0.001; \*\*\*\* - <0.0001.

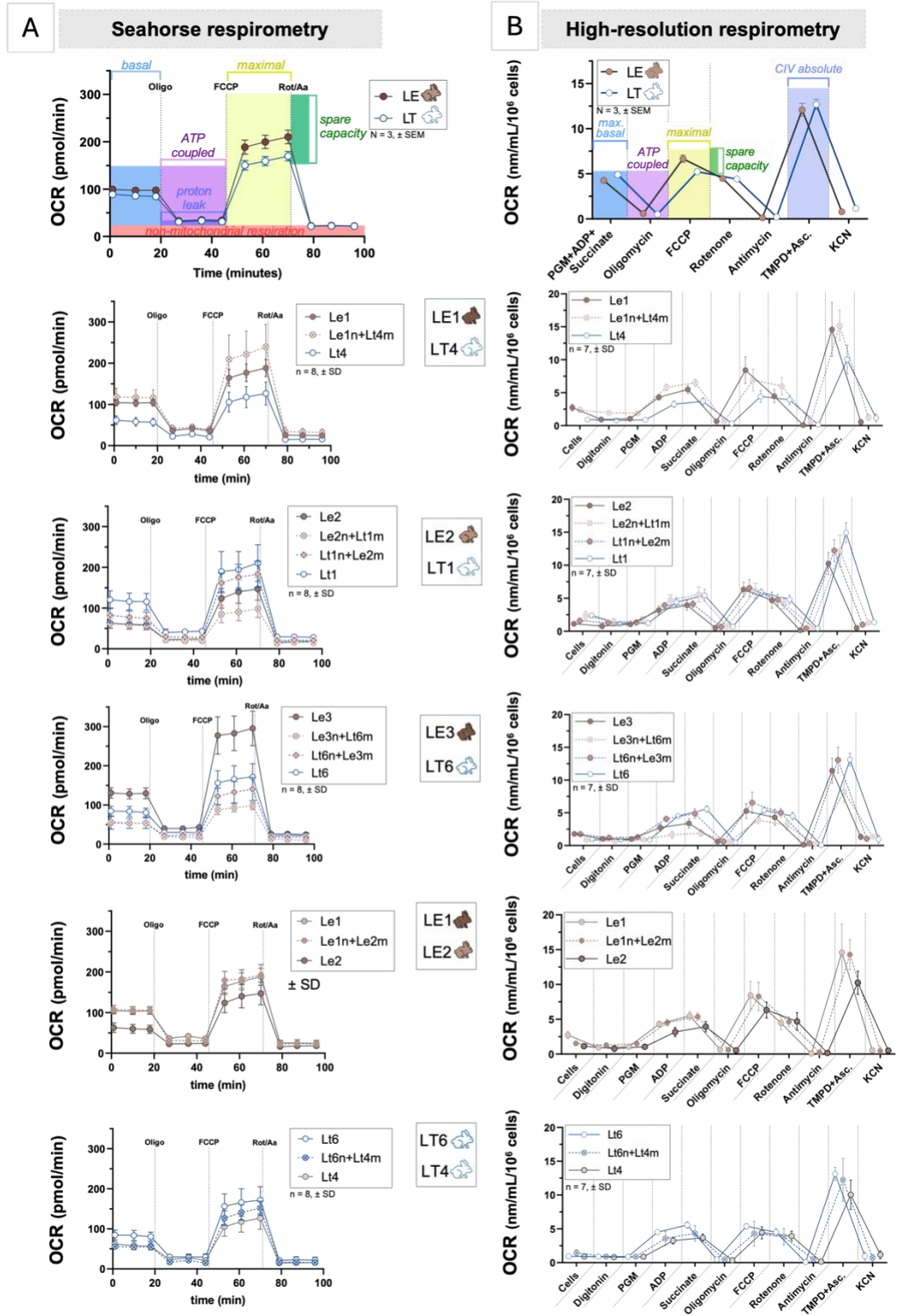

**Supplementary Fig. 4.** Respiration traces of all studied cell lines measured (A) in intact cell using Seahorse Cell Mito Stress assay and (B) in permeabilized cells using high-resolution respirometry. OCR, oxygen consumption rate. n, nucleus; m, mtDNA. Data presented as mean ± SD. *p*-value: \* - $<0.05$; \*\* - $<0.01$; \*\*\* - $<0.001$; \*\*\*\* - $<0.0001$.

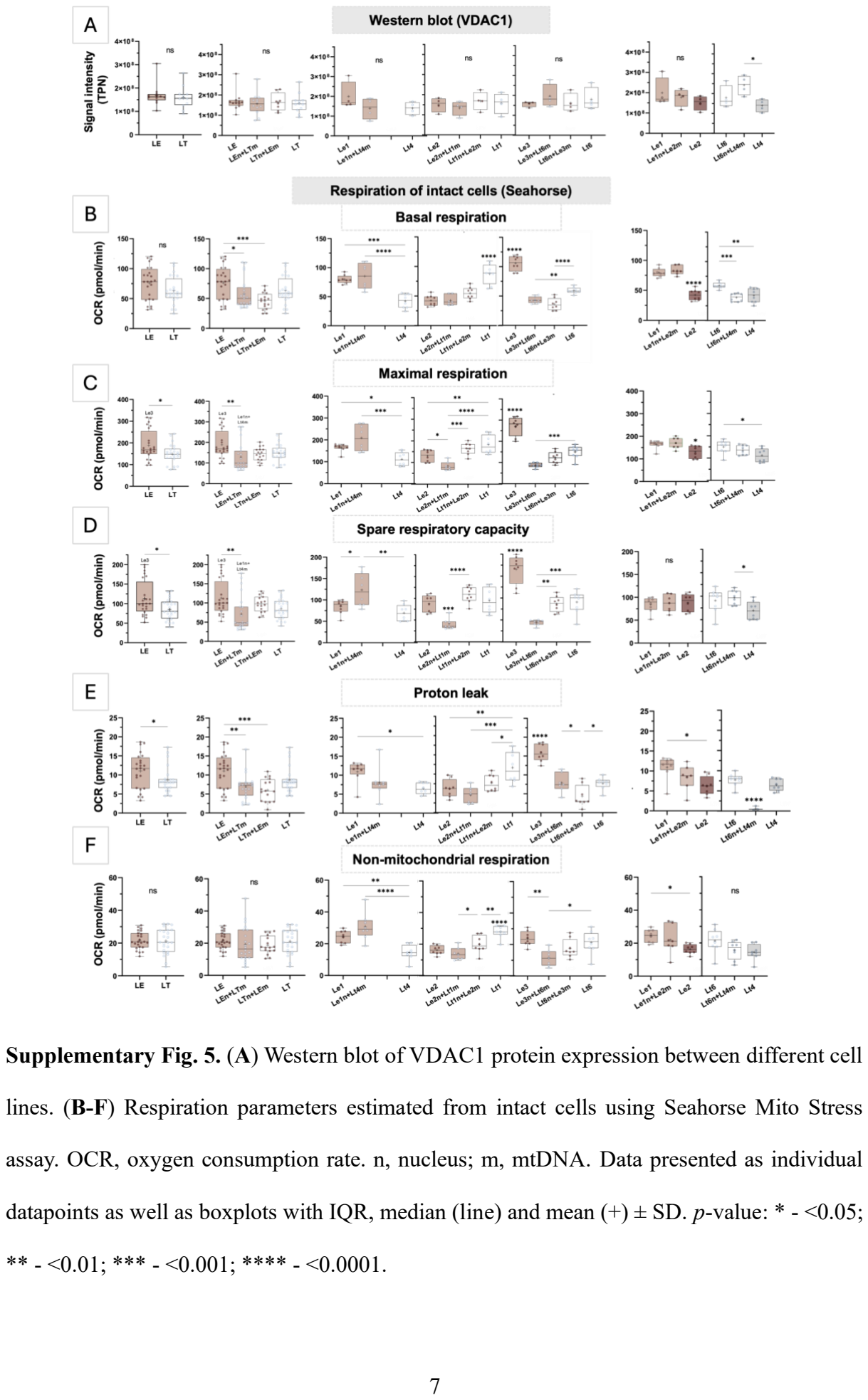

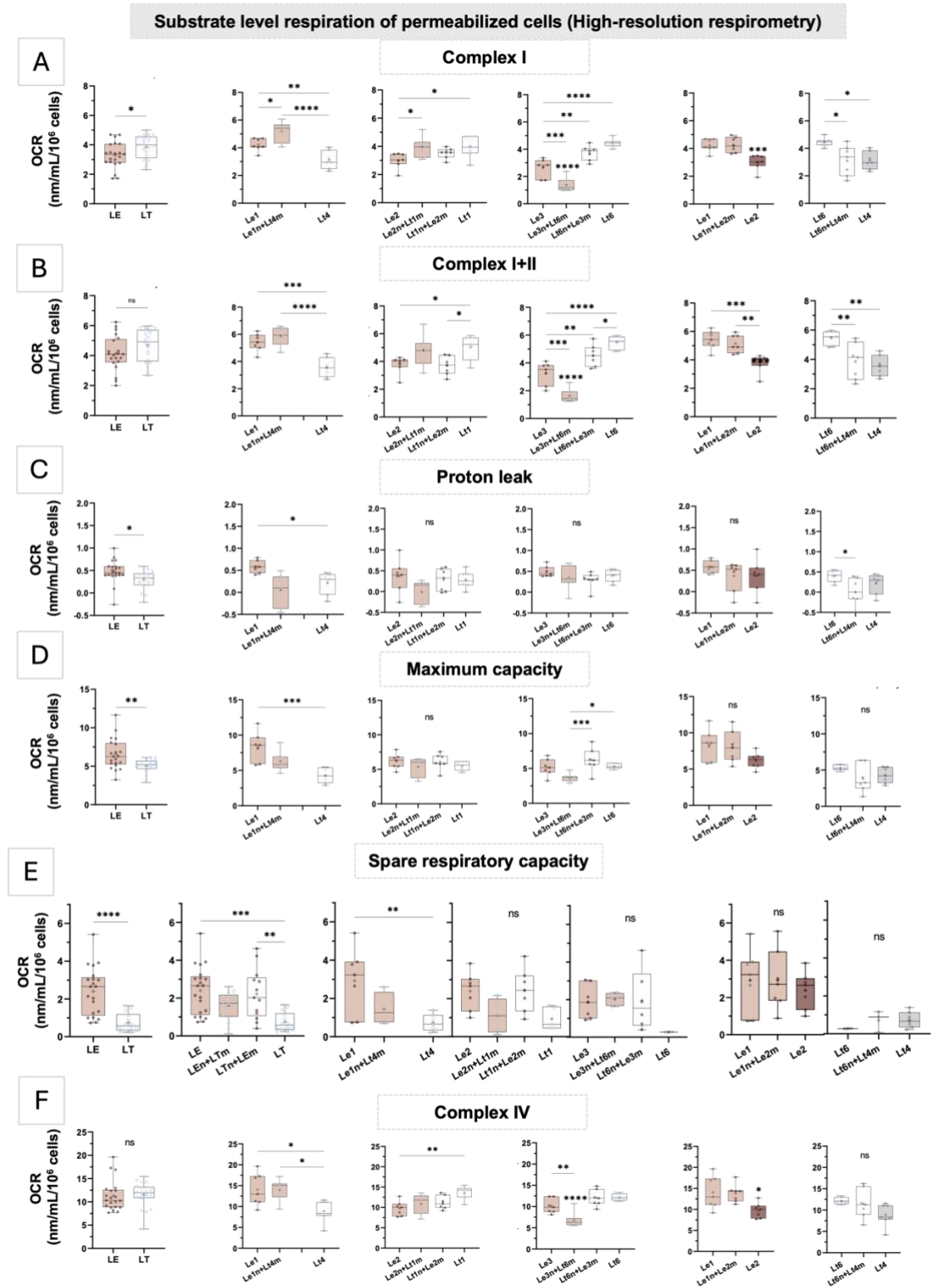

**Supplementary Fig. 6. (A-F)** Respiration parameters estimated from permeabilized cells using high-resolution respirometry. OCR, oxygen consumption rate. n, nucleus; m, mtDNA. Data presented as individual datapoints as well as boxplots with IQR, median (line) and mean (+) ± SD. *p*-value: \* - <0.05; \*\* - <0.01; \*\*\* - <0.001; \*\*\*\* - <0.0001.

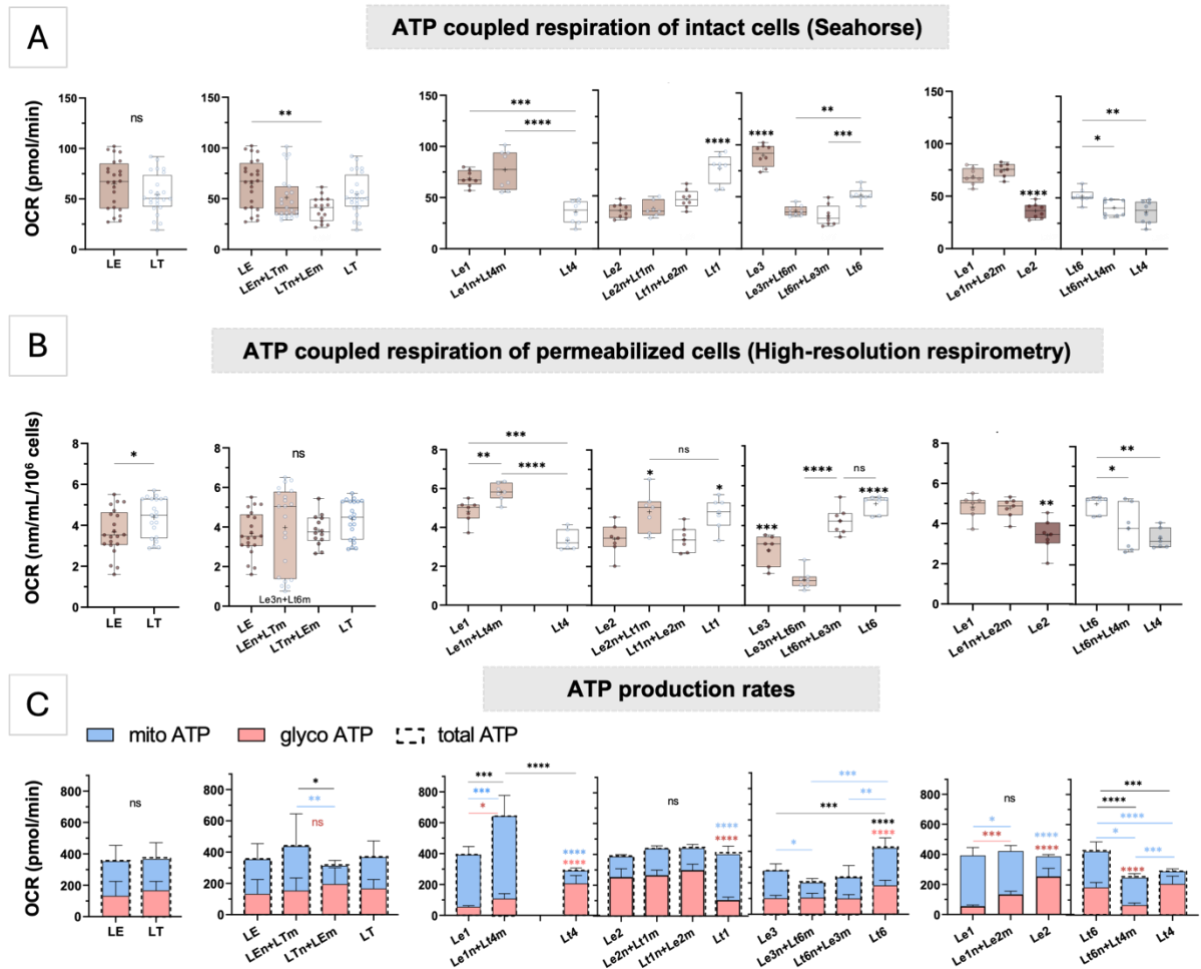

**Supplementary Fig. 7.** ATP coupled respiration measured (A) in intact cell using Seahorse Cell Mito Stress assay and (B) in permeabilized cells using high-resolution respirometry. (C) ATP production rates estimated with Seahorse Real-Time ATP Rate assay. Bar charts show oxidative (mitochondrial) in blue, glycolytic in red and total ATP production rates measured in all cell lines. OCR, oxygen consumption rate. n, nucleus; m, mtDNA. Data presented as individual datapoints as well as boxplots with IQR, median (line) and mean (+) ± SD. *p*-value: \* - $<0.05$; \*\* - $<0.01$; \*\*\* - $<0.001$; \*\*\*\* - $<0.0001$.

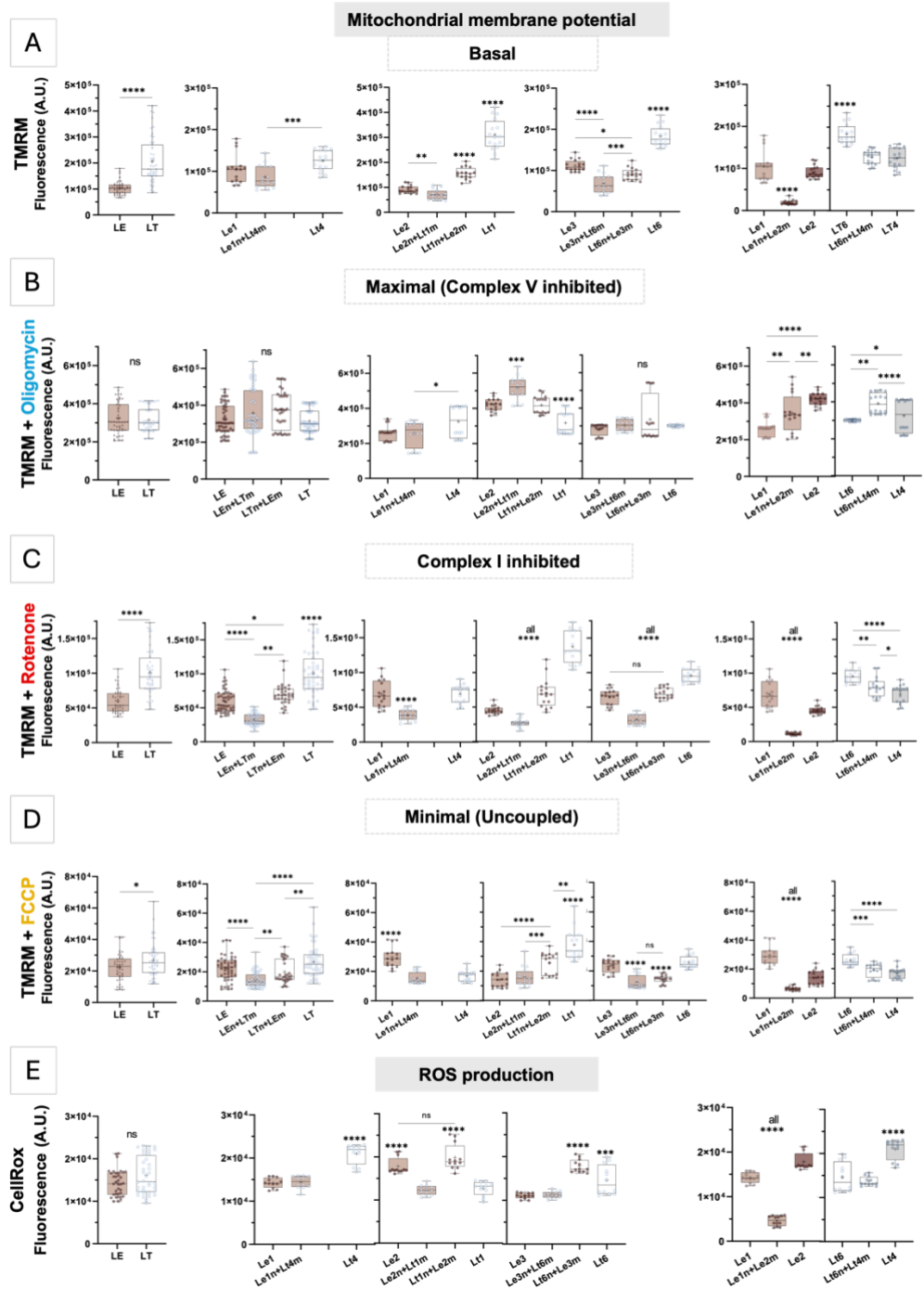

**Supplementary Fig. 8.** (A-D) Mitochondrial membrane potential estimated with TMRM staining (E) Cellular reactive oxygen species (ROS) production estimated with CellRox staining. n, nucleus; m, mtDNA. Data presented as individual datapoints as well as boxplots with IQR, median (line) and mean (+) ± SD. *p*-value: \* - $<0.05$; \*\* - $<0.01$; \*\*\* - $<0.001$; \*\*\*\* - $<0.0001$.

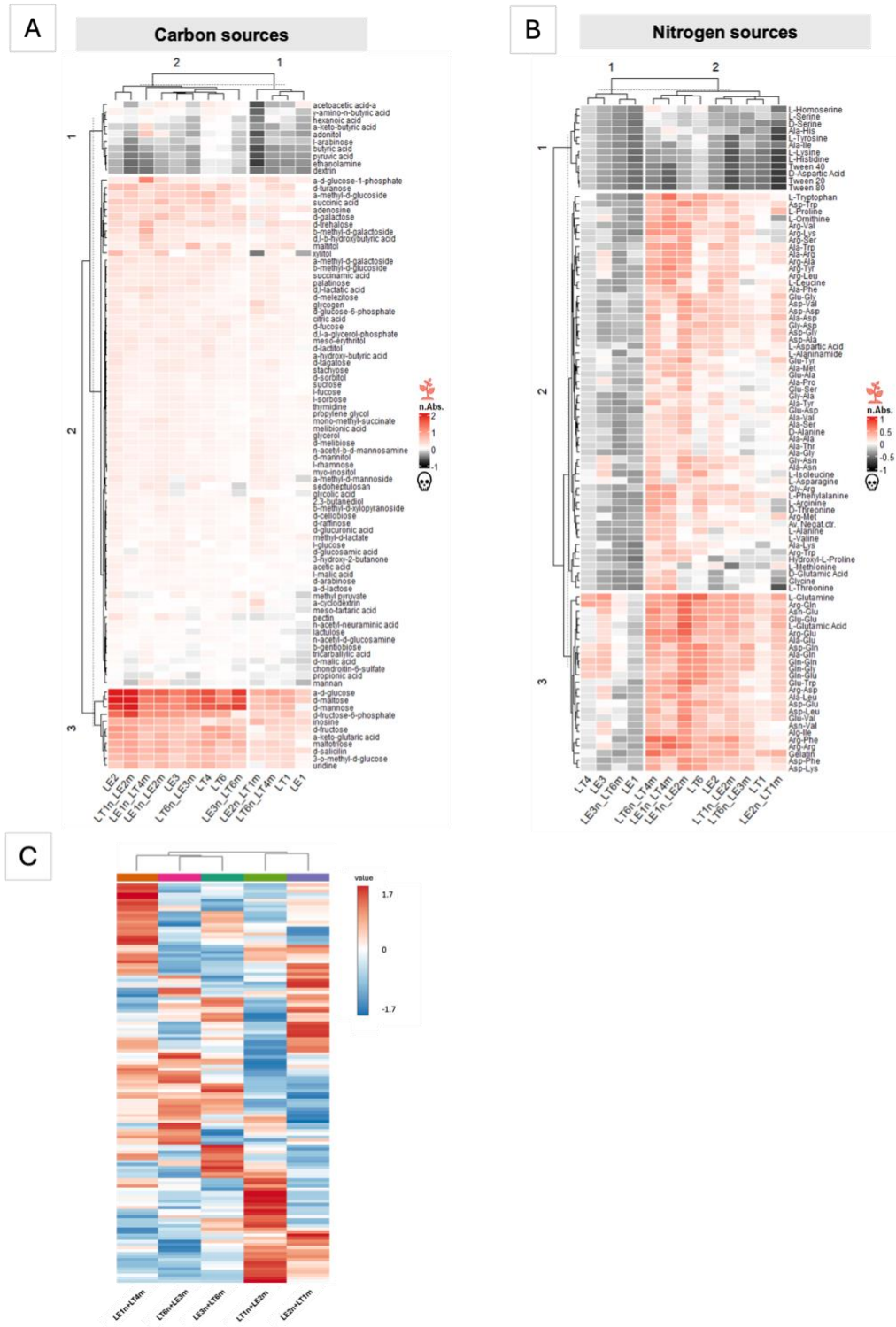

**Supplementary Fig. 9.** (A) Heatmaps of Biolog phenotype assay of carbon substrates and (B) nitrogen substrates. Cell proliferation and viability were supported by preferred substrates (in red) and depleted by disliked substrates (in black). (C) Clustering of the heterocybrids based on all 130 identified metabolites. n, nucleus; m, mtDNA.

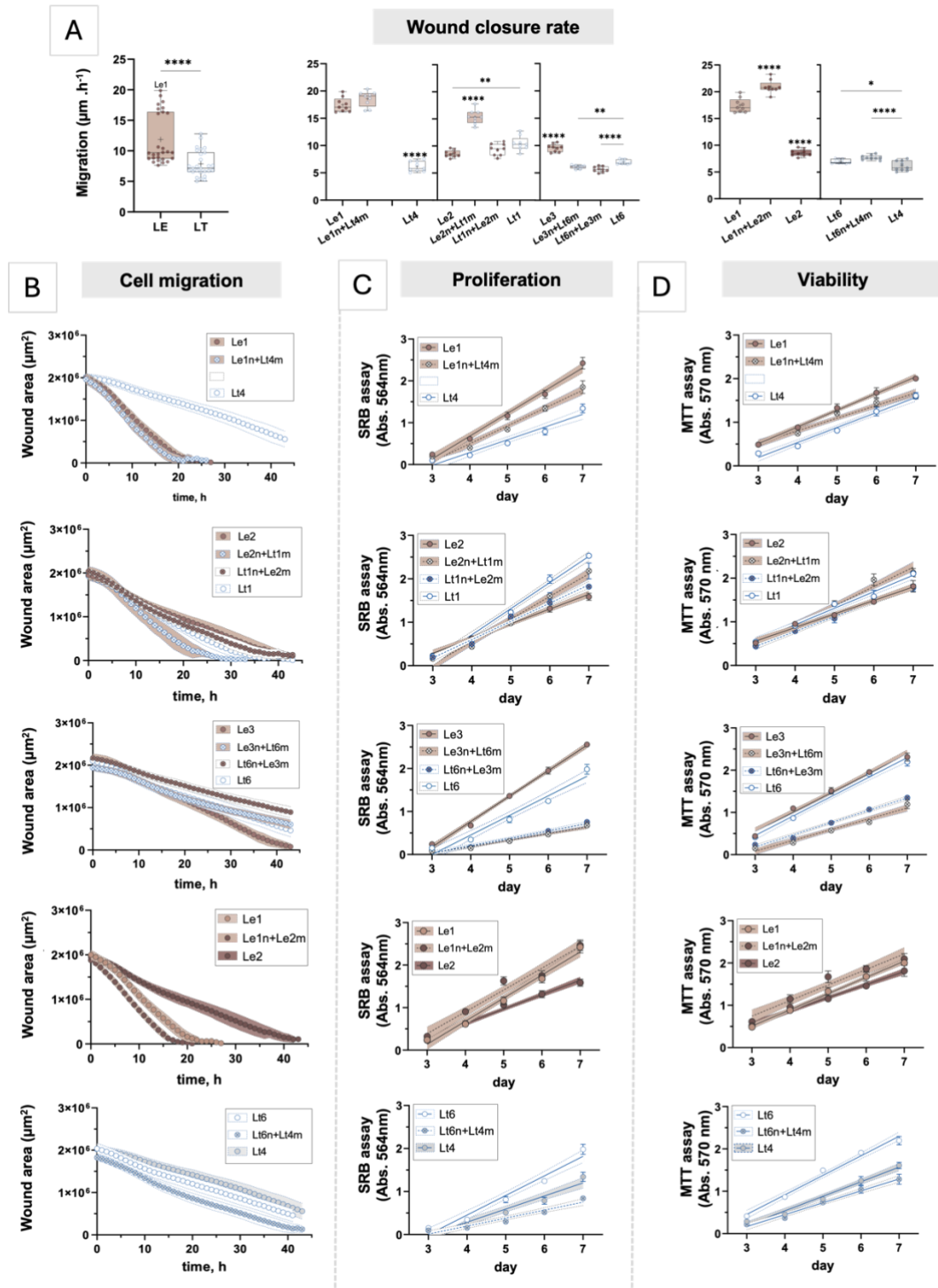

**Supplementary Fig. 10.** (A) Wound closure rate estimated from (B) Cell migration recorded in wound healing assay. Data presented as individual datapoints as well as boxplots with IQR, median (line) and mean (+) ± SD. (C) Cell proliferation. (D) Cell viability. n, nucleus; m, mtDNA. Data presented as mean ± SD. *p*-value: \* - $<0.05$; \*\* - $<0.01$; \*\*\* - $<0.001$; \*\*\*\* - $<0.0001$.

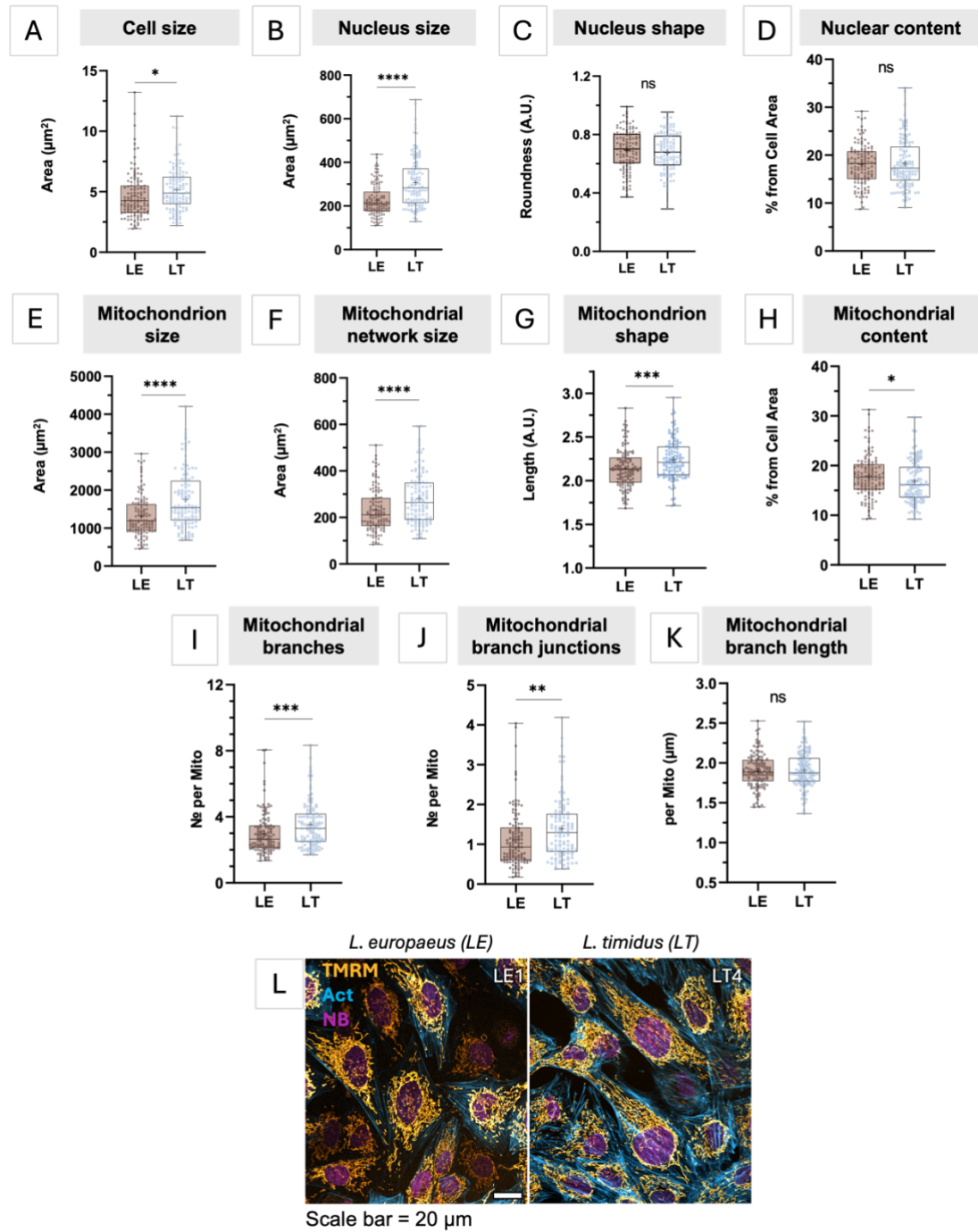

**Supplementary Fig. 11. (A-K)** Comparison of cellular morphometric parameters between LE and LT fibroblasts. N = 3. Data presented as individual datapoints as well as boxplots with IQR, median (line) and mean (+) ± SD. $p$-value: \* - $<0.05$; \*\* - $<0.01$; \*\*\* - $<0.001$; \*\*\*\* - $<0.0001$. **(L)** Representative microscopy images of hare fibroblasts. Nuclei were stained with NucBlue (NB) shown in purple, mitochondria with TMRM shown in yellow and cytoskeleton with CellMask Actin (Act) shown in blue. Scale bar = 20 $\mu\text{m}$.

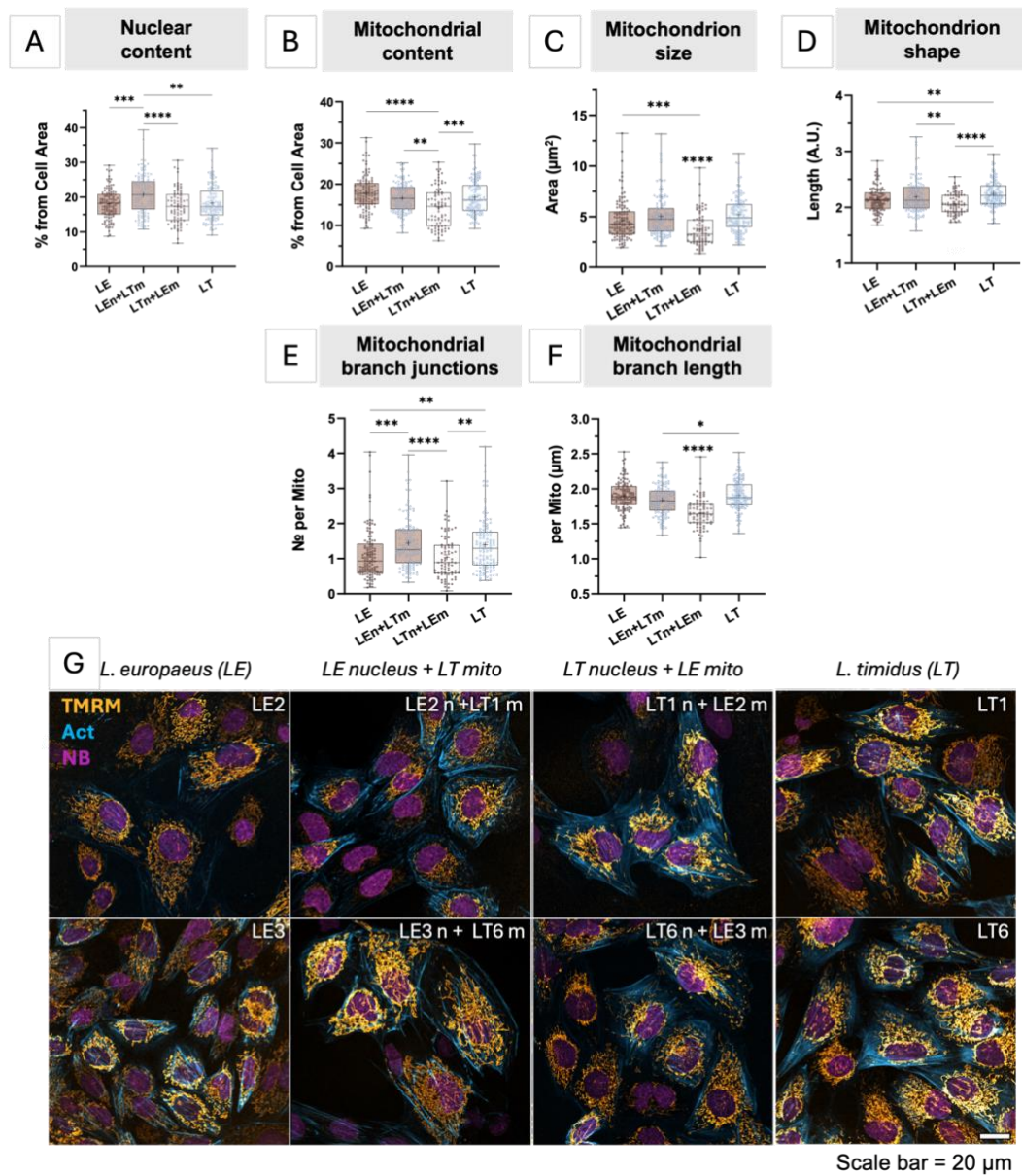

**Supplementary Fig. 12.** (A-F) General comparison of cellular morphometric parameters between heterocybrids, LE and LT parental fibroblasts. N = 3, however false LT4n+LE1m heterocybrid has been removed from analyses. Data presented as individual datapoints as well as boxplots with IQR, median (line) and mean (+) ± SD. *p*-value: \* - <0.05; \*\* - <0.01; \*\*\* - <0.001; \*\*\*\* - <0.0001. (L) Representative microscopy images of heterocybrids and their parental cell lines. Nuclei were stained with NucBlue (NB) shown in purple, mitochondria with TMRM shown in yellow and cytoskeleton with CellMask Actin (Act) shown in blue. Scale bar = 20 μm. n, nucleus; m, mtDNA.

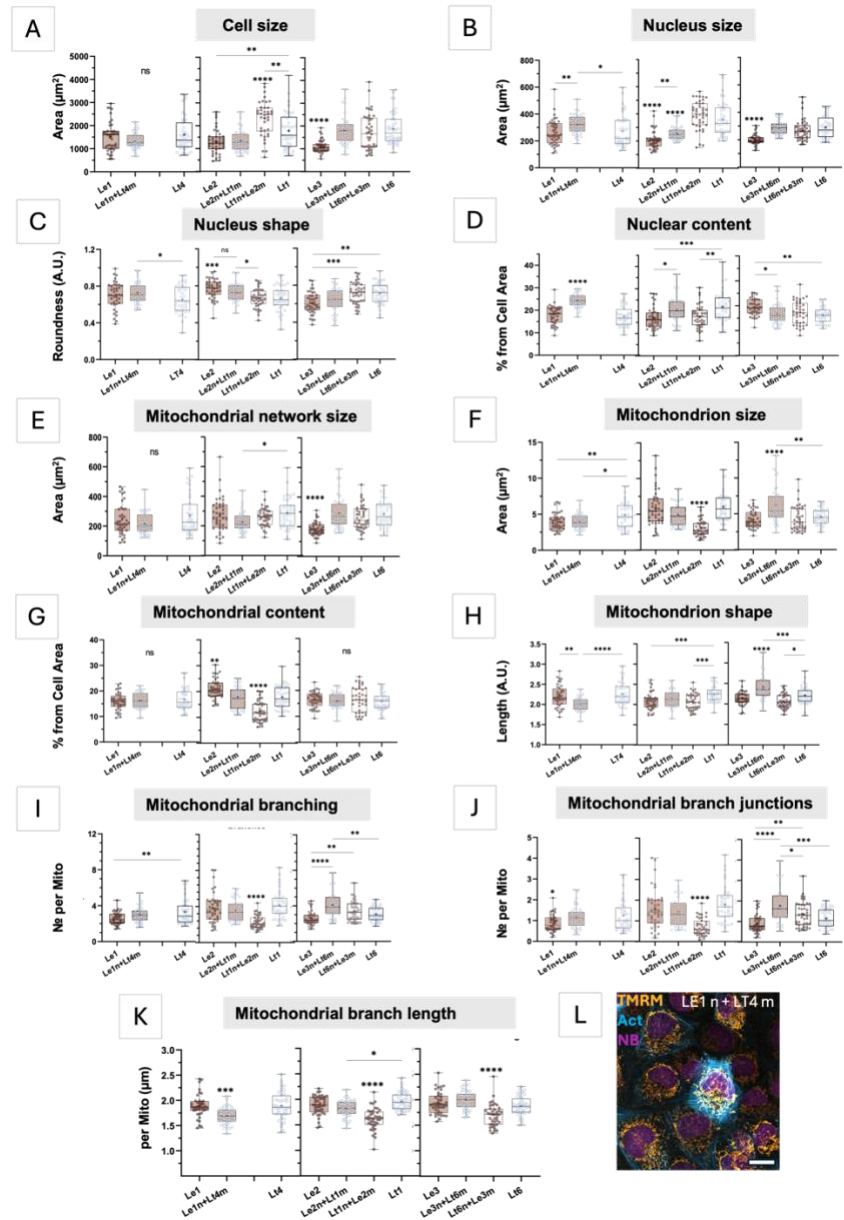

**Supplementary Fig. 13. (A-K)** Comparison of cellular morphometric parameters between heterocybrids and their parental cell lines. False LT4n+LE1m heterocybrid has been removed from analyses. Data presented as individual datapoints as well as boxplots with IQR, median (line) and mean (+) ± SD. *p*-value: \* - <0.05; \*\* - <0.01; \*\*\* - <0.001; \*\*\*\* - <0.0001. **(L)** Representative microscopy image of the LE1n+LT4m heterocybrid cell line. Nuclei were stained with NucBlue (NB) shown in purple, mitochondria with TMRM shown in yellow and cytoskeleton with CellMask Actin (Act) shown in blue. Scale bar = 20 $\mu\text{m}$. n, nucleus; m, mtDNA.

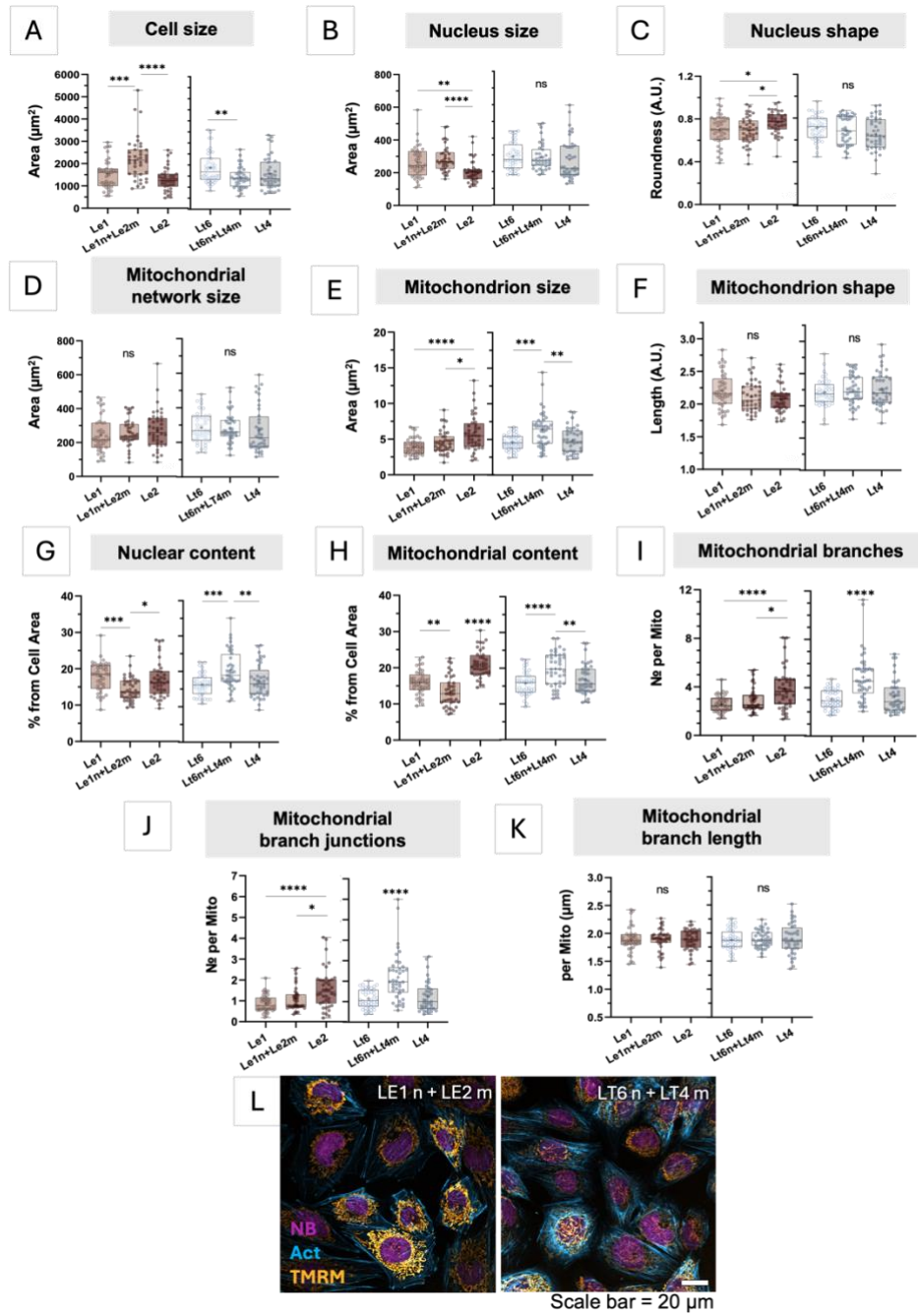

**Supplementary Fig. 14.** (A-K) Comparison of cellular morphometric parameters between control homocybrids and their parental cell lines. Data presented as individual datapoints as well as boxplots with IQR, median (line) and mean (+) ± SD. *p*-value: \* - $<0.05$; \*\* - $<0.01$; \*\*\* - $<0.001$; \*\*\*\* - $<0.0001$. (L) Representative microscopy images of control homocybrids. Nuclei were stained with NucBlue (NB) shown in purple, mitochondria with TMRM shown in yellow and cytoskeleton with CellMask Actin (Act) shown in blue. Scale bar = 20 $\mu\text{m}$. n, nucleus; m, mtDNA.
